# Dodecyl Creatine Ester Improves Cognitive Function and Identifies Drivers of Creatine Deficiency

**DOI:** 10.1101/2022.11.03.514982

**Authors:** Aloïse Mabondzo, Rania Harati, Léa Broca-Brisson, Anne-Cécile Guyot, Narciso Costa, Francesco Cacciante, Elena Putignano, Laura Baroncelli, Matthew R Skelton, Cathy Saab, Emmanuelle Martini, Henri Benech, Thomas Joudinaud, Jean-Charles Gaillard, Jean Armengaud, Rifat A. Hamoudi

## Abstract

Creatine transporter deficiency prevents creatine uptake into the brain, leading to mental retardation. To better understand the pathophysiology, this study focuses on the identification of biomarkers related to cognitive improvement in a Slc6a8 knockout mouse model (Slc6a8/y) engineered to mimic the clinical features of CTD patients which have low brain creatine content. Shotgun proteomics analysis of 4,035 proteins in four different brain regions; the cerebellum, cortex, hippocampus (associated with cognitive functions) and brain stem, and muscle as a control, was performed in 24 mice. Comparisons of the protein abundance in the four brain regions between DCE-treated intranasally Slc6a8-/y mice and wild type and DCE-treated Slc6a8-/y and vehicle group identified 14 biomarkers, shedding light on the mechanism of action of DCE. Integrative bioinformatics and statistical modeling identified key proteins associated with CTD, including KIF1A and PLCB1. The abundance of these proteins in the four brain regions was significantly correlated with both the object recognition and the Y-maze tests. Functional analysis confirmed their key roles and associated molecules in CTD pathogenesis.

## Introduction

Neurodevelopmental disorders represent a significant health problem due to the heterogeneity of the underlying causes and a lack of appropriate treatment options. Creatine transporter deficiency (CTD) is a rare genetic disorder and a subset of intellectual disability (ID). However its symptoms including autism-like symptoms with ID, expressive speech and language delay, movement disorders, and epilepsy (DesRoches *et al*, 2015; Salomons *et al*, 2001; Stockler *et al*, 2007) overlaps with those of common neurodevelopmental disorders. CTD is an X-linked (XL) disorder caused by mutations in *SLC6A8*, the gene encoding creatine transporter (CrT)(Braissant *et al*, 2011), that prevent the transport of creatine (Cr), which is essential for brain function, into the brain (van de Kamp *et al*, 2014) and is estimated to be the cause of 1-2% of all cases of XL ID (DesRoches *et al*., 2015) (Cheillan *et al*, 2012) and about 1% of cases with ID of unknown aethiology (Clark *et al*, 2006). As expected, the symptoms are most severe in males, with female carriers presenting with a milder phenotype. Some of the main difficulties in elucidating the pathogenesis of and treating ID are that there are wide variety of causes of ID, with no single cause being associated with a significant majority of ID cases.

Several combinations of nutritional supplements or Cr precursors l-arginine and l-glycine, have been studied as therapeutic approaches for CTD, but they have shown very limited success (Bruun *et al*, 2018; Valayannopoulos *et al*, 2013)’ (Jaggumantri *et al*, 2015). However, our previous findings suggest that following the inhalation of dodecyl creatine ester (DCE) in Slc6a8^-/v^ (CrT KO) mice, an animal model recapitulating the clinical features of human CTD, an increase of Cr brain content and synaptic markers could be achieved in the synapsis terminals and thus improving the cognitive function of Slc6a8-/y mice. These findings highlight that DCE might be a therapeutic option for CTD patients (Trotier-Faurion *et al*, 2013; Trotier-Faurion *et al*, 2015; Ullio-Gamboa *et al*, 2019a).

To gain a better understanding of the pathophysiology of CTD and DCE treatment efficacy, we focused in this study on the identification of biomarkers related to cognitive improvement in a Slc6a8-/y mice which have low brain Cr content. To that end, the study employed a combination of cognitive tests and molecular methods to decipher some of the molecular mechanisms involved in CTD pathophysiology. The cognitive tests included the use of object recognition test (ORT), Y-maze and Morris water maze (MWM) tests to show the decline of cognitive function in *Slc6a8^-/v^* mice and whether the treatment with DCE improves the cognitive function. The molecular methods involved the application of shotgun proteomics to four different brain regions; the cerebellum, cortex, hippocampus (associated with cognitive functions) and brain stem, and muscle as a control.

## Results

### Restoration of cognitive functions in DCE-treated creatine knockout mice

DCE was intranasally administered as previously reported (Ullio-Gamboa *et al*, 2019b) to CrT KO mice for 30 days, while wild-type (WT) and vehicle-treated mice were used as controls (N=8 per group). A volume of 6 μL of DCE or vehicle was placed in the nostril. DCE (4 mg/g) or vehicle was given twice bilaterally (12 μL total volume). Consistent with previous findings, CrT KO mice showed object recognition deficits when compared with WT mice. Mice were assigned to treatment groups by sorting animals based on discrimination index (DI) and alternating assignments between vehicle and treatment to avoid performance confounds.

Fig. 1a shows DI data. Vehicle-treated CrT KO mice showed reductions in DI compared with both WT mice (p=0.023) and DCE-treated CrT KO mice (p < 0.05). DCE-treated CrT KO mice spent more time exploring the novel object than vehicle-treated CrT KO mice (one-way ANOVA, p < 0.01; Tukey’s post hoc test, p < 0.05; Fig. 1b), but there was no difference between DCE-treated CrT KO mice and WT mice (p = 0.406). Strikingly, the median exploration time for the DCE-treated CrT KO mice was 96% of the median exploration time for the WT mice.

**Fig. 1.**
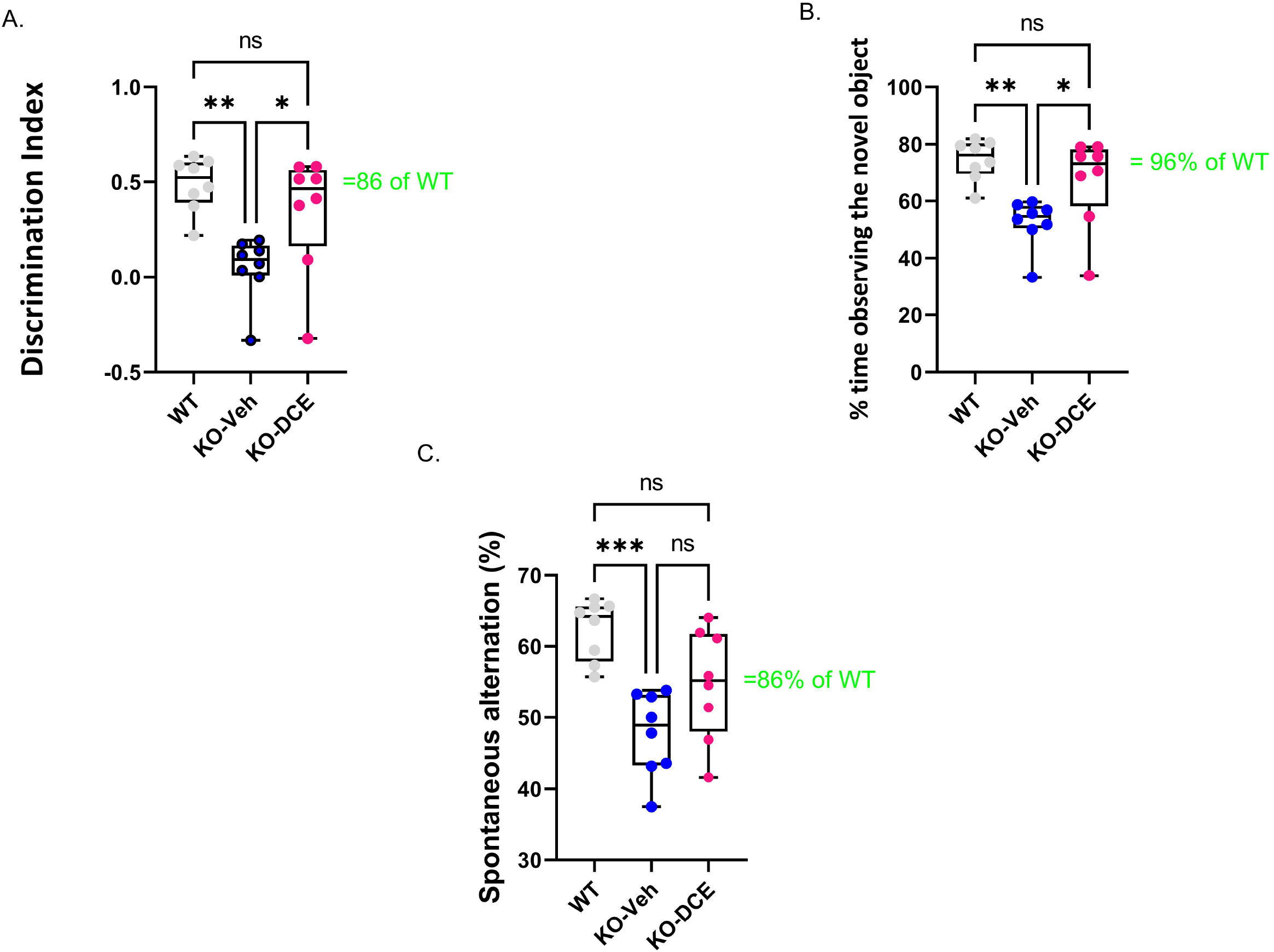
Analysis of cognition among the different experimental groups. **a-b,** Object recognition in the ORT was impaired in CrT KO mice. a, Analysis of the object discrimination index and b, percentage of time spent exploring the novel object revealed that WT and DCE-treated CrT KO mice but not vehicle-treated CrT KO mice showed a preference for the novel object. **c,** Early deficiency of working and spatial memory in CrT KO mice measured by the Y-maze test. CrT KO mice showed a change in the spontaneous alternation percentage in the Y-maze test, which was significantly improved by DCE treatment. The data are the mean ± s.e.m. Statistical analysis was performed by one-way ANOVA followed by Tukey’s post-hoc test, *p < 0.05; **p < 0.001; ns= not significant.

We used the Y-maze test measuring the spontaneous alternation rate, the single parameter with the highest accuracy in discriminating WT and CrT KO mice (Raffaele *et al*, 2020). A reduction of the spontaneous alternation rate in vehicle-treated CrT KO mice (48%) was observed compared to WT mice (62%; one-way ANOVA, p < 0.001; Tukey’s post hoc test, p < 0.001; Fig. 1c). The median spontaneous alternation rate was higher in DCE-treated CrT KO mice than in vehicle-treated CrT KO mice but did not differ between DCE-treated CrT KO mice and WT mice (86%; p = 0.058; Fig. 1c).

Memory was assessed using the MWM test. The results show that DCE did not have a beneficial effect on the performance of CrT KO mice in the MWM test, suggesting that DCE treatment ameliorates some cognitive deficits seen in these mice.

### Proteomic analysis identified unique proteins abundant in specific brain regions

In order to identify proteins across the different brain regions involved in the pathogenesis of CTD, proteomics based differential abundance analysis was performed in the different brain regions of 24 mice. Immediately after behavioral testing, the animals from all three groups were sacrificed. Label-free shotgun proteomics analysis of cortical, hippocampal, cerebellar, and brainstem tissues and muscle tissue as a control was carried out for each mouse in WT, vehicle-treated CrT KO mice and DCE-treated CrT KO mice. High-resolution tandem mass spectrometry (MS) analysis of the 120 biological samples generated a large dataset comprising of 7,006,153 MS/MS spectra showing the abundance of 4,035 proteins for the five tissues from each mouse. Following unsupervised filtering and normalization (Hachim *et al*, 2020) (Supplemental Table 1), the proteins whose abundance was significantly altered by mutant CrT and DCE treatment were identified using reproducibility-optimized statistical testing in each of the five tissue types. The workflow of the bioinformatics analysis is described in Supplemental Fig. 1.

A total of 376, 322, 163, and 321 proteins were found to be differentially abundant in the cortex, cerebellum, brainstem and hippocampus, respectively, between WT mice and vehicle-treated CrT KO mice. The abundances of 320, 416, 279, and 323 proteins in the cortex, cerebellum, brainstem and hippocampus, respectively, were marked altered in DCE-treated CrT KO mice compared with vehicle-treated CrT KO mice (Supplemental Fig. 2a-d; Supplemental Fig. 3b, d, f, h), while 413, 223, 176 and 175 proteins were differentially abundant in DCE-treated CrT KO mice compared with WT mice (Supplemental Fig. 3a, c, e, g). Overlapping proteins were observed in the comparisons of DCE-treated vs. vehicle-treated mice CrT KO, DCE-treated CrT KO vs. WT mice, and vehicle-treated CrT KO vs. WT mice. Most of the overlapping proteins in the comparison between DCE-treated vs. vehicle-treated mice CrT KO and vehicle-treated CrT KO vs. WT mice were found in the cortex (98 proteins) (Fig. 2a), cerebellum (153 proteins) (Fig. 2b), and hippocampus (126 proteins) (Fig. 2c); conversely, fewer overlapping proteins were found in the brainstem (53 proteins) (Fig. 2d; Supplemental Table 2). The proteins whose abundances were altered in vehicle-treated mice and DCE-treated mice compared with WT mice were also selected in cortex (11 proteins) (Fig. 2a), cerebellum (7 proteins) (Fig. 2b), hippocampus (10 proteins) (Fig. 2c) and brain stem (1 protein) (Fig. 2d; Supplemental Table 2).

**Fig. 2.**
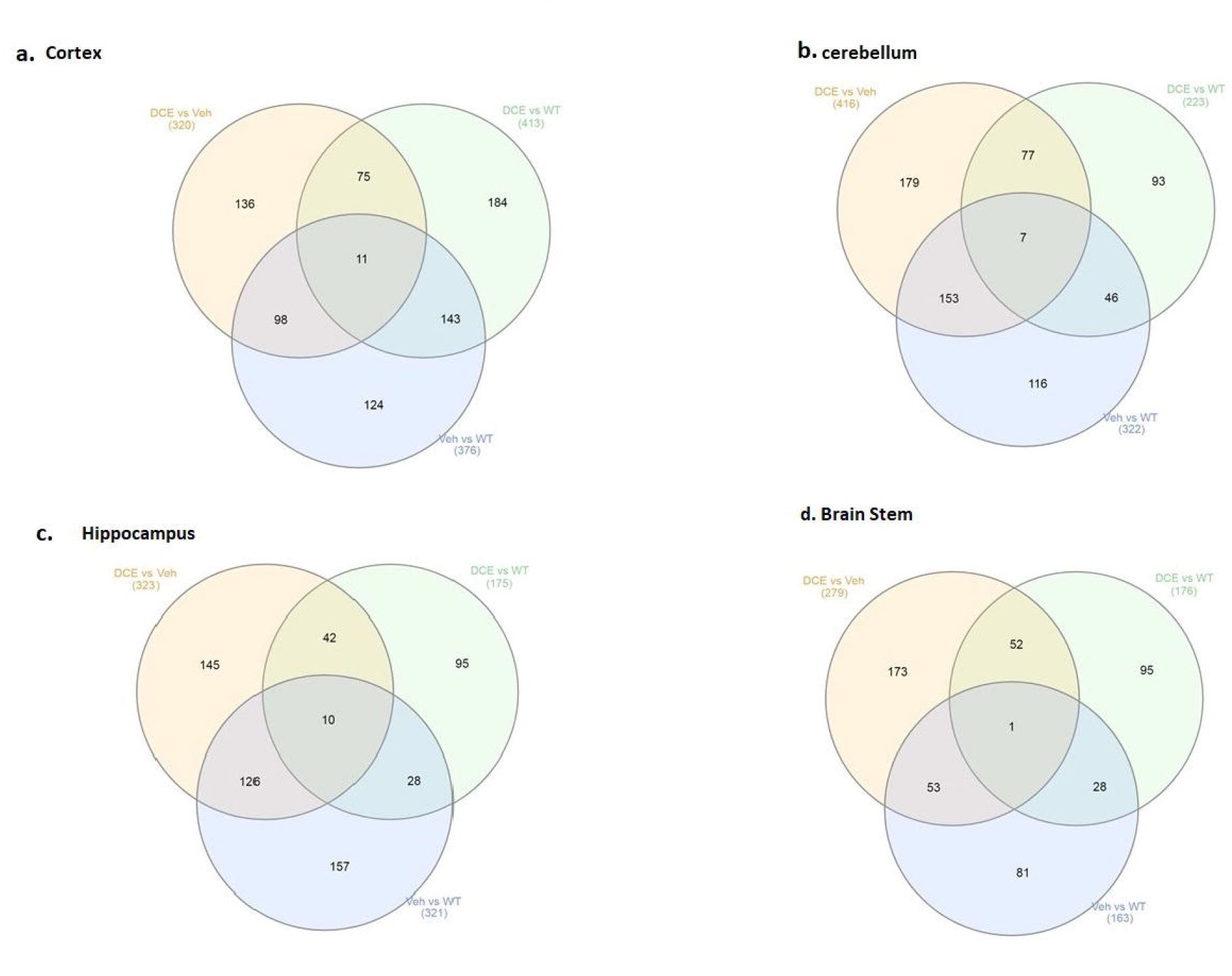
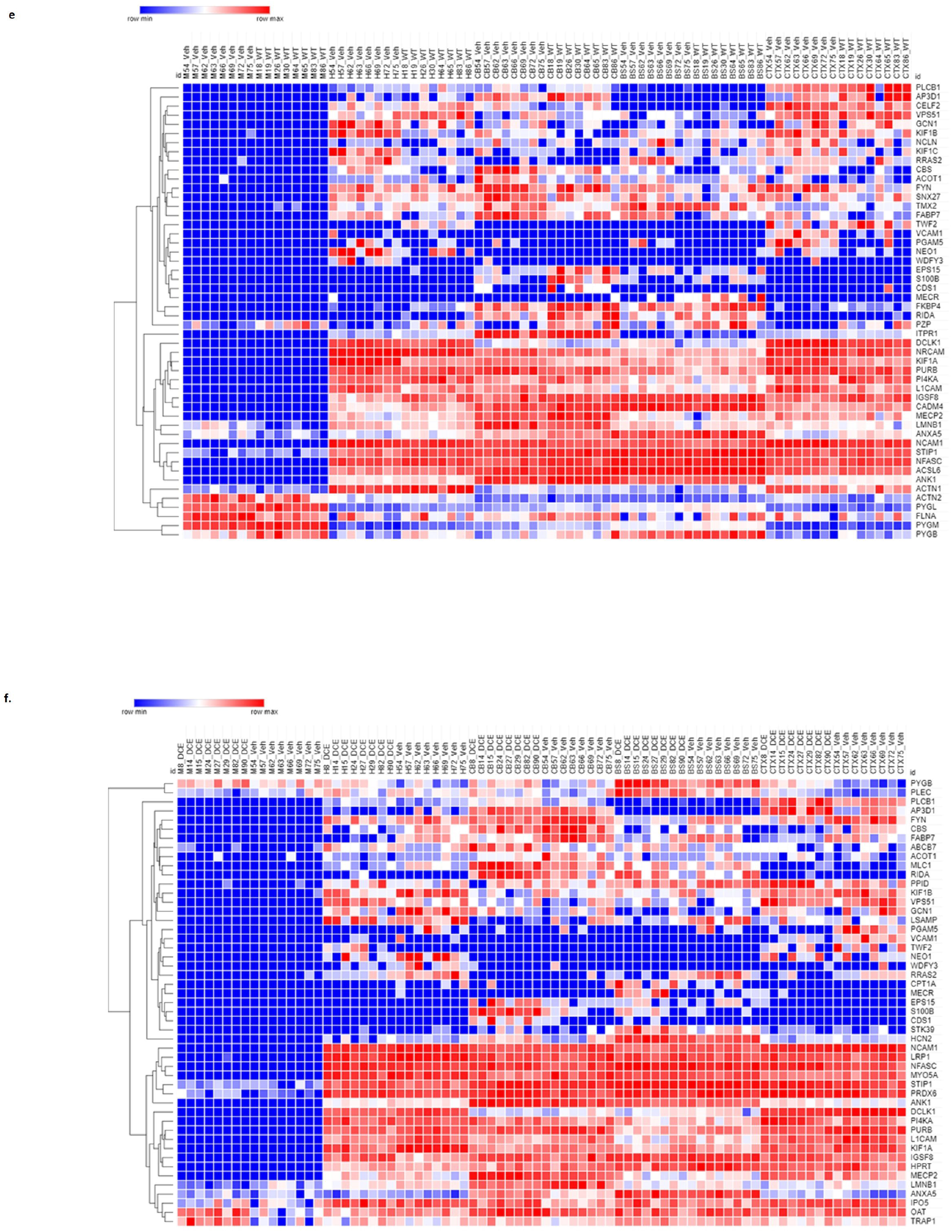
Comparison of proteomic signatures in different brain regions among the different experimental groups. **a-d,** Venn diagram showing overlapping proteins among the three experimental groups. The proteins that showed a significant change in abundance in CrT KO mice compared with the WT and in DCE-treated mice compared with vehicle-treated mice were selected for pathway analysis. The proteins that showed a change in abundance in the vehicle-treated mice and the DCE-treated mice compared with the WT mice were also selected for pathway analysis. The abundances of these proteins in the cortex, cerebellum, brainstem, hippocampus and muscle were analyzed using a multivariate statistical model based on one-way ANOVA followed by Bonferroni’s post hoc test. **e,** Heatmap showing the proteins that showed a significant change in abundance in vehicle-treated CrT KO mice compared WT mice. **f**, Heatmap showing the proteins that showed a significant change in abundance in DCE-treated CrT KO mice compared with vehicle-treated CrT KO mice.

The overlapping proteins whose abundances were significantly altered in all CrT KO mice compared to that in WT mice and in DCE-treated mice CrT KO mice compared to that in vehicle-treated CrT KO mice were then selected for pathway analysis using gene set enrichment analysis carried out using ENRICHR. The proteins found to be involved in different diseases and pathways were selected for subsequent analysis (Cortex (41 proteins), Hippocampus (53 proteins), Cerebellum (52 proteins), and brain stem (25 proteins). These findings suggest that lack of Cr into the brain of CrT KO mice leads to a significant alteration of protein abundance involved in the pathogenesis of CTD.

### Identification of a panel of 14 proteins in the brain involved in cognitive activity using a multivariate statistical model

A multivariate statistical model comparing WT mice with vehicle-treated CrT KO mice (Fig. 2e) and DCE-treated CrT KO mice with vehicle-treated CrT KO mice was used to identify key proteins (Fig. 2f). Among the proteins whose abundances were affected by either CrT deficiency or DCE treatment, that were discussed in the previous section, bioinformatics analysis and statistical modelling identified 14 proteins that were the most abundant in the four different brain regions in both vehicle-treated and DCE-treated CrT KO mice. Those most abundant proteins were significantly altered by the mutation compared to WT(vehicle-treated vs WT) and by the treatment compared to vehicle (DCE-treated vs. vehicle treated) and their abundance after treatment was restored to levels comparable to those in WT mice (Fig. 3a; Supplemental Tables 3 & 4).

**Fig. 3.**
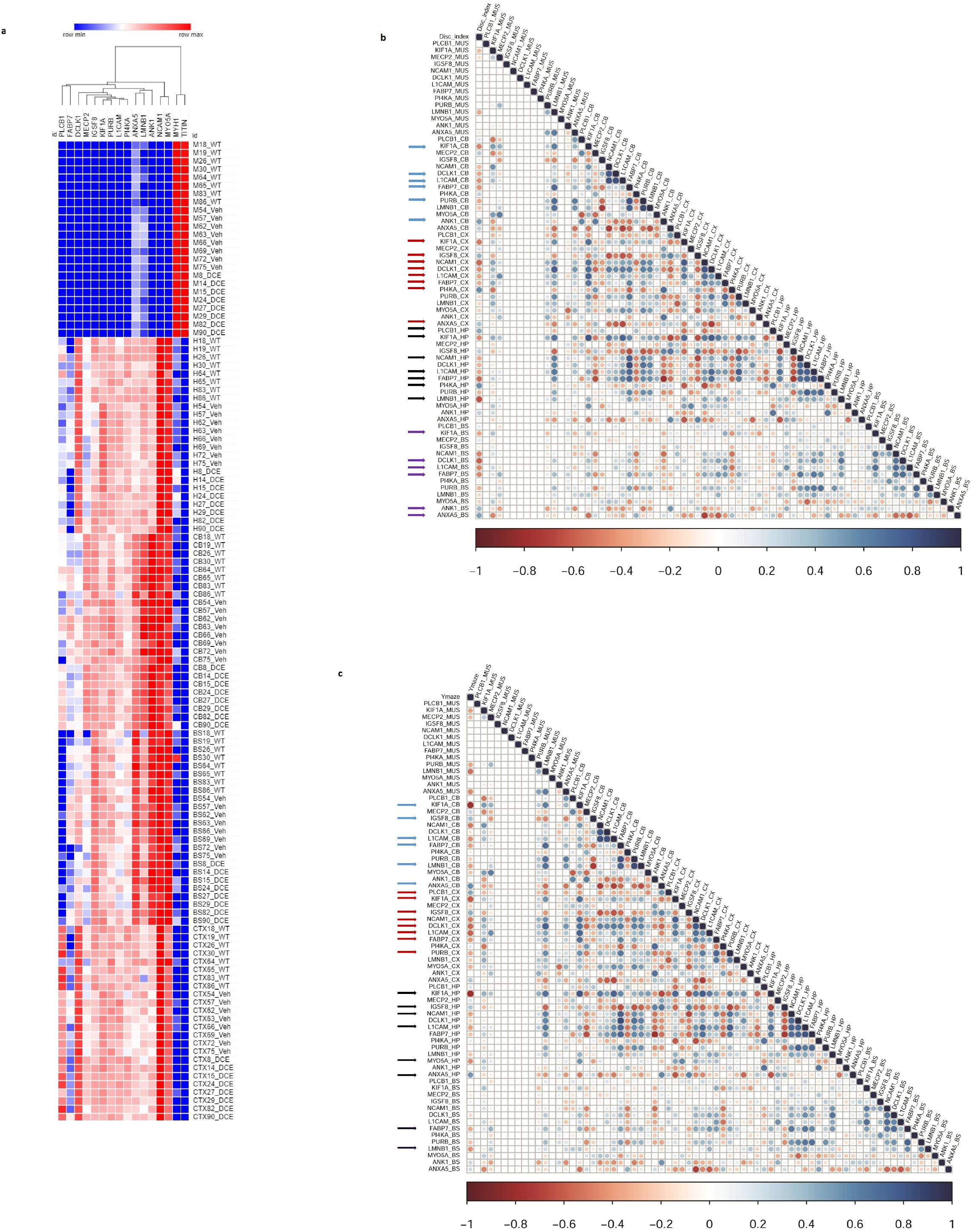
Correlation of the levels of 14 proteins with performance in different cognitive tests using a stepwise regression statistical model. **a,** Heatmap showing the abundance levels of the 14 proteins that showed significant changes in abundance in both vehicle-treated CrT KO mice and DCE-treated CrT KO mice. **b**, Map showing the correlation between the level of each of the 14 proteins with performance in the ORT to evaluate cognition. The arrows indicate correlations between the levels of specific proteins and performance in the ORT. **c,** Maps showing the correlation between the level of each of the 14 proteins and performance in the Y-maze test. The arrows indicate correlations between the levels of specific proteins and performance in the Y-maze test. The correlation maps were derived using a stepwise regression model to assess the correlation between the abundance each of the 14 differentially expressed proteins and performance in each of the cognitive tests. Group comparisons were carried out using a statistical model based on one-way ANOVA followed by Bonferroni’s post hoc test for multiple comparisons.

The results showed that PLCB1 abundance in the cortex (p=0.0003) (Supplemental Table 3a); MeCP2 (p=0.022), ANK1 (p=0.0001), and ANXA5 (p=0.00004) abundance in the cerebellum (Supplemental Table 3c); and IGSF8 (p=0.00036) abundance in the hippocampus (Supplemental Table 3b) were significantly downregulated in vehicle-treated CrT KO mice compared with WT mice, DCE treatment significantly rescued the abundance of these proteins in these brain regions in CrT KO mice (Fig. 3a; Supplemental Table 4). In contrast, NCAM1 (p=1.38E-07), PI4KA (p=0.026), DCLK1 (p=3^-15^) and PURB (p=0.045) abundance in the cortex (Supplemental Table 3a); KIF1A (p=2^-24^), NCAM1 (p=0.038) and L1CAM (p=0.001) abundance in the hippocampus (Supplemental Table 3b); and LMNB1 (p=1E^-05^), FABP7 (p= 8.30E^-05^), and PURB (p=0.03) abundance in the cerebellum (Supplemental Table 3c) were significantly upregulated in CrT-treated vehicle KO mice compared with WT mice but were not different between WT and DCE-treated CrT KO mice (Fig. 3a; Supplemental Table 4).

### Correlation of the proteins with cognitive behavioral tests using stepwise regression statistical model shows KIF1A is abundant across the brain regions

A stepwise regression model was used to assess the effect of each of the proteins of interest on cognitive outcomes to identify those that may influence cognitive function. The levels of several proteins were significantly correlated with the discrimination index (DI); the correlation between KIF1A, Fabp7 and L1CAM levels and DI was found in the hippocampus, cortex, cerebellum and brain stem (n= 11, Supplemental Table 5a).

We also observed that the level of LMNB1, which is involved in cerebellar ataxia and adult autosomal dominant leukodystrophy (r^2^ = −0.634, p < 0.0001), and the level of Pi4KA (r^2^= −0.634, p= 0.066) in the hippocampus were correlated with the DI. Several proteins were also significantly correlated with the performance in the Y-maze test (n= 11, Fig. 3c, Supplemental Table 5b). The level of KIF1A, a kinesin that transports synaptic vesicle precursors, was most strongly correlated with the spontaneous alternation rate in the Y-maze test (hippocampus: r^2^ = −0.795, p<0.0001; cortex: r^2^ = −0.608, p= 0.001; cerebellum: r^2^ = 0.774, p < 0.0001). Fabp7 and L1CAM correlated with the spontaneous alternation rate in the Y-maze test in the 3 brain regions (Supplemental Table 5b). In addition, PLCB1 is significantly more abundant in the cortex compared to other brain regions and its abundance in the cortex and hippocampus is correlated with DI and Y-maze (p = 0.01).

Amongst the 14 key proteins with individual animal performance in different cognitive tests, KIF1A was shown to be the protein with the most uniformly abundance across the 4 brain regions and correlated with the DI in the novel object recognition test and spontaneous alternation in the Y-maze test.

### KIF1A and PLCB1 interplay is associated with DCE treatment efficacy in CrT KO mice

KIF1A participates in vesicular transport, and vesicles containing the neurotrophin BDNF have been found to be under the control of KIF1A (Kondo *et al*, 2012). BDNF is produced in the neocortex throughout brain development and accelerates the overall redistribution of cortical neurons (Iki *et al*, 2005). DCE-mediated rescue of KIF1A levels in the cortex (Fig. 4a), hippocampus (Fig. 4b), and cerebellum (Fig. 4c) of CrT KO mice resulted in higher levels of pro-BDNF/BDNF (p=0.0043; Fig. 4d) a long-term potentiation (LTP) biomarkers, which are linked to cognitive function improvement. These results indicate that KIF1A is a potential key player in CTD pathogenesis.

**Fig. 4.**
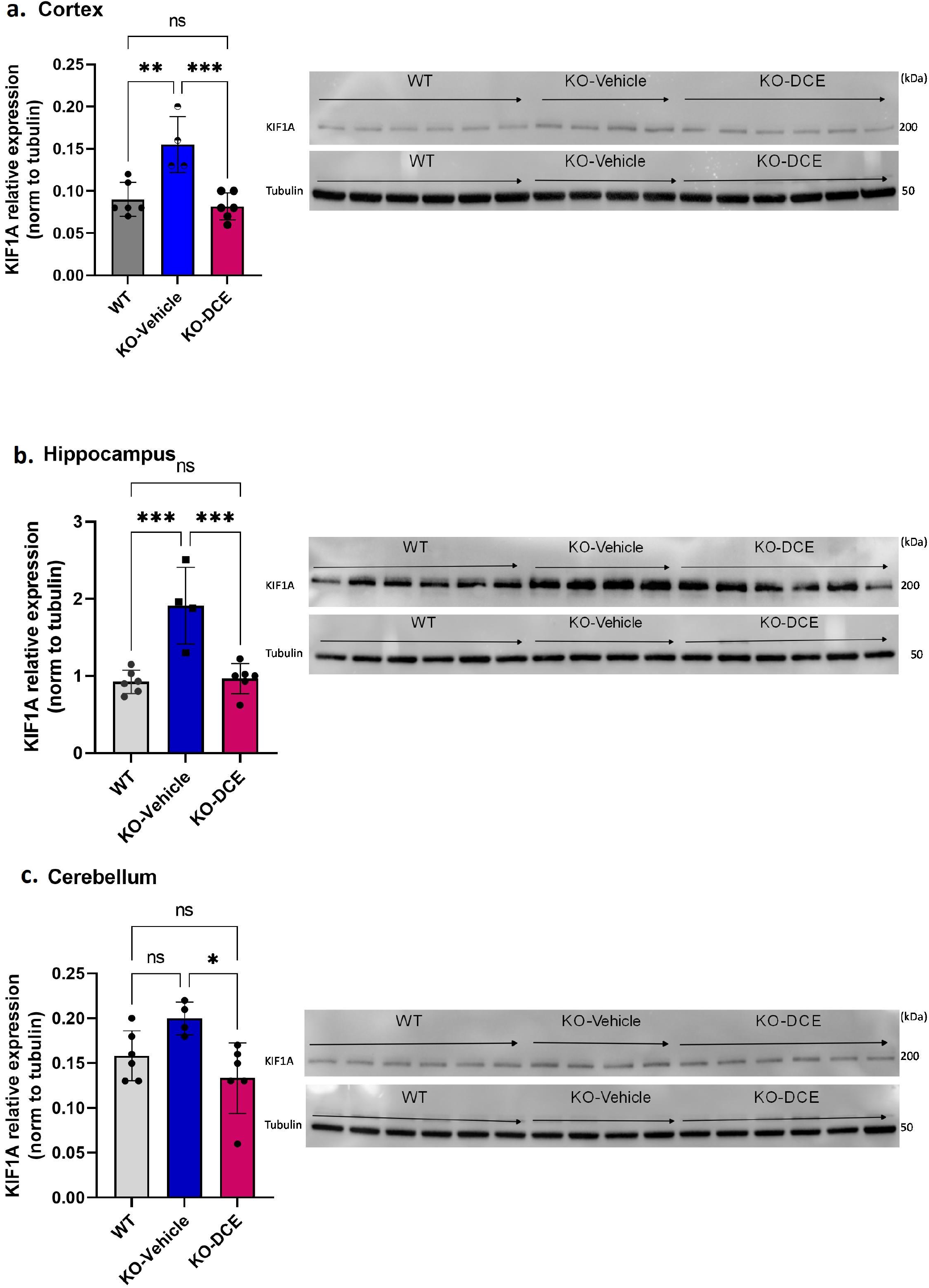
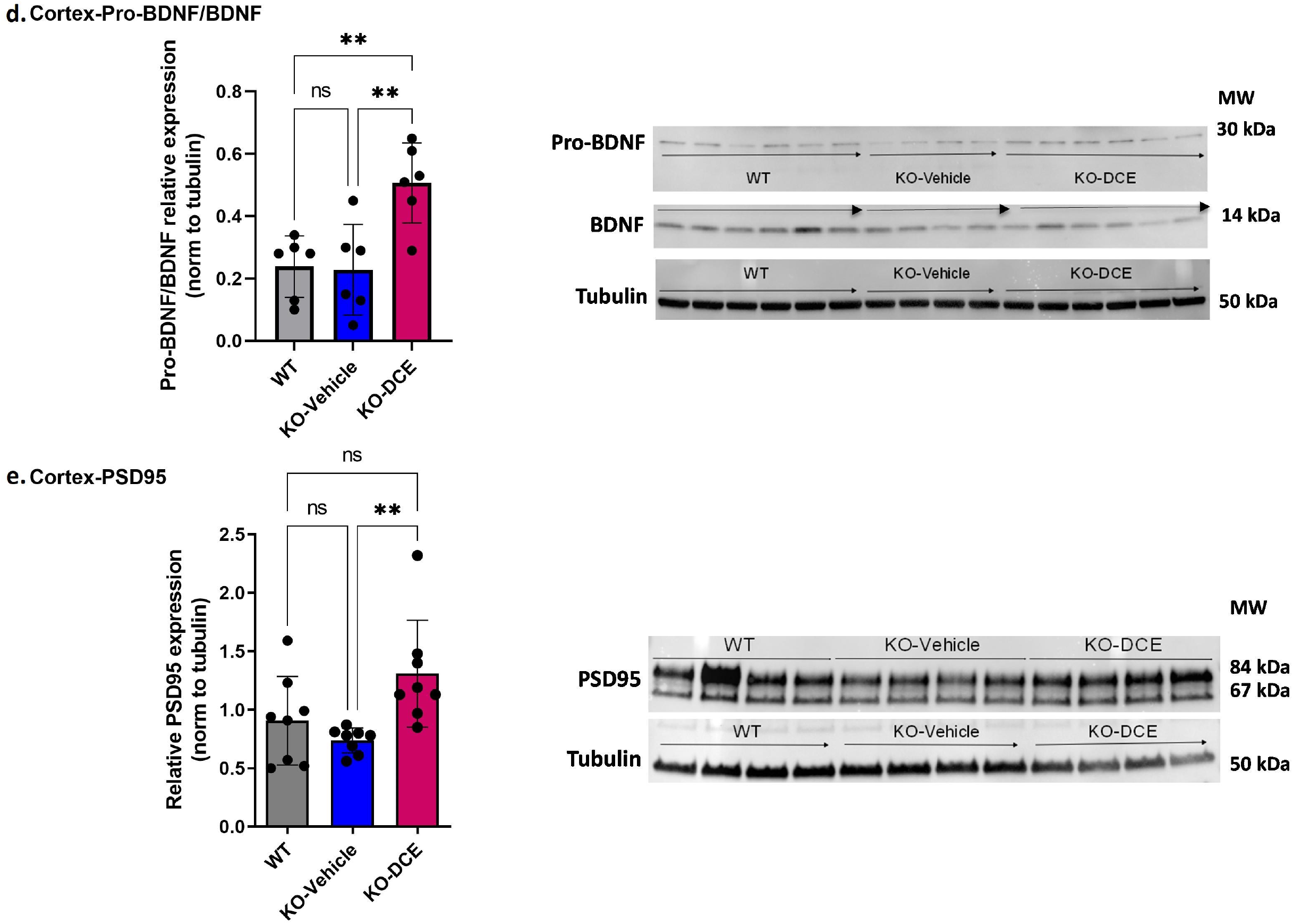
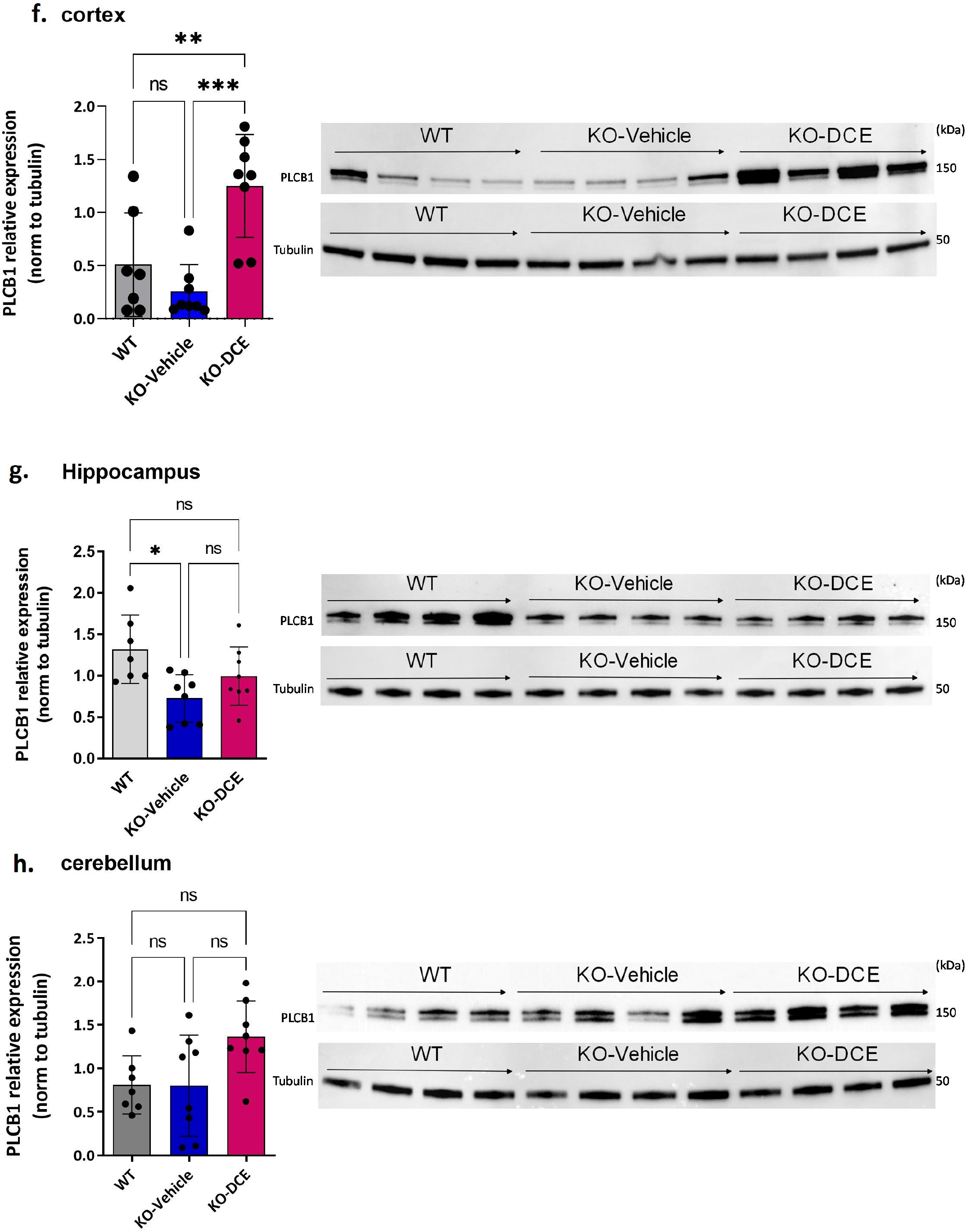
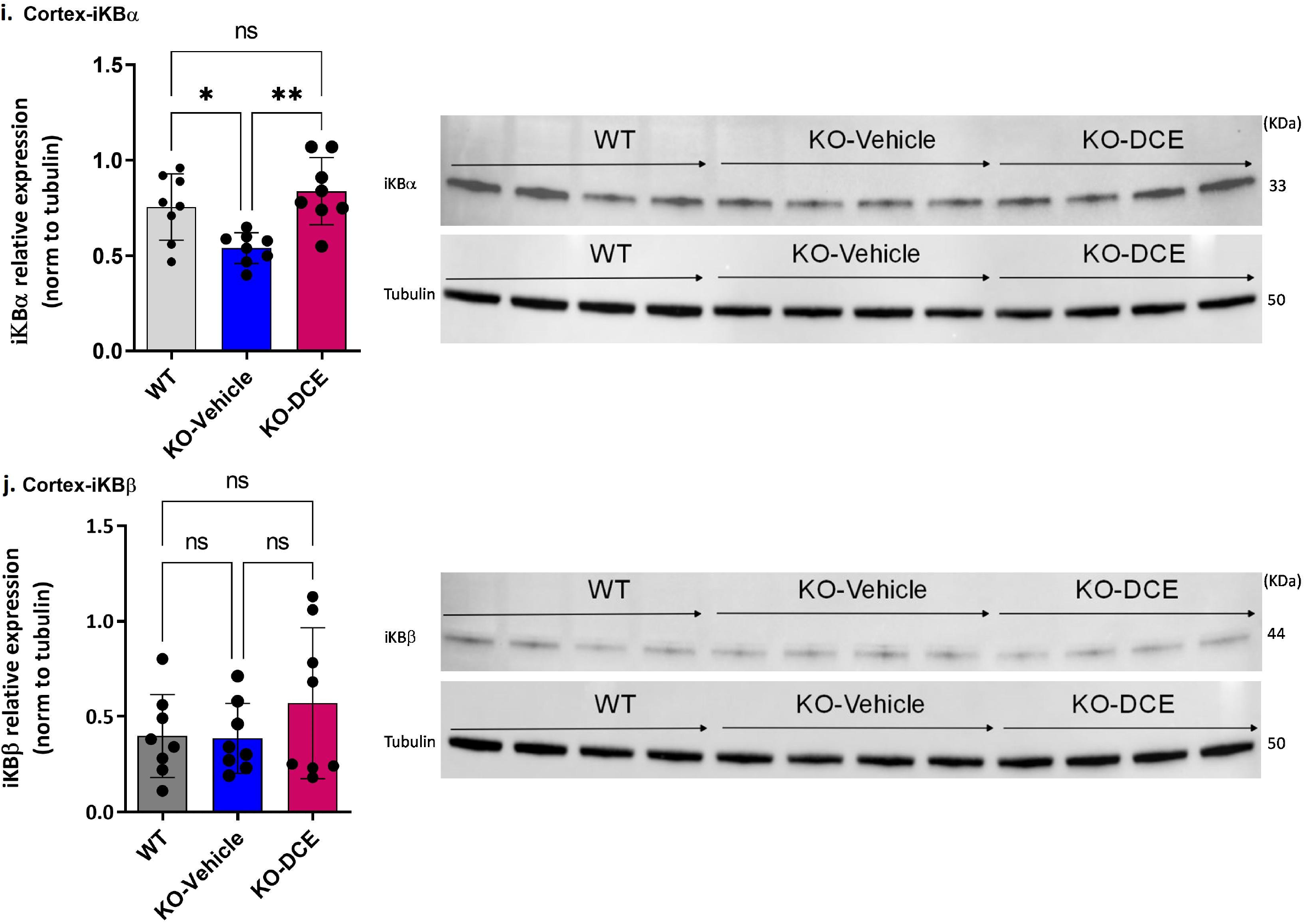
KIF1A and PLCB1 interplay is associated with DCE treatment efficacy in CrT Ko mice., **a-c,** Western blot results showing that the KIF1A level was significantly increased in the cortex (**a**), hippocampus (b) and cerebellum (c) in vehicle-treated CrT KO mice compared to WT mice, while DCE treatment rescued this overabundance in the three brain regions in CrT KO mice. **d, e,** Western blot results showing that the pro-BDNF/BDNF ratio (d) was significantly altered in DCE-treated CrT KO mice compared to that of both WT mice and vehicle-treated CrT KO mice. In addition, the PSD95 level (**e**) was significantly altered in the cortices of vehicle-treated CrT KO mice compared to that of DCE-treated CrT KO mice, f-h, Western blotting showed that PLCB1 protein (150 kDa) abundance was increased in the cortex (**f**), hippocampus (g) and cerebellum (**h**) in CrT KO mice at 0 days after DCE treatment. **i, j**, Western blot results for Ibα (**i**) and IκBβ (**j**) showing that DCE promotes IκBα transcription factor abundance but not IκBβ transcription factor abundance. The data are the mean ± s.e.m. (n=8/group). Statistical analysis was performed by one-way ANOVA followed by Tukey’s post-hoc test. Tubulin was used as a loading control. *p≤0.05; **p≤0.001; ***p≤0.0001; ns= not significant.

KIF1A overabundance in the developing brains of CrT KO mice likely leads to synaptic dysfunction, thus contributing to cognitive and memory impairments (Iki *et al*., 2005; Lu *et al*, 2014). We found that in the cortex, the abundances of the presynaptic protein IgSF8 protein (Fig. 3a), which has been reported to be a critical regulator of brain microcircuits and neuronal function (Apostolo *et al*, 2020), and the postsynaptic density protein PSD95 were altered in vehicle-treated CrT KO mice compared with the abundances in WT mice (Fig. 4e). These results indicate that KIF1A is a potential key player in CTD pathogenesis.

In order to highlight the central role of KIF1A abundance across the different brain regions of CrT Ko mice, proteins co-immunoprecipitated together with KIF1A were identified by MS. Shotgun proteomics revealed that KIF1A and PLCB1 could be co-immunoprecipitated with anti-KIF1A antibody from cortical and hippocampal extracts. In addition, PLCB1 abundance was correlated with the DI in the object recognition test (Supplemental Table 5a) and spontaneous alternation rate in the Y-maze test (Supplemental Table 5b) (hippocampus: r2 = 0.460, p=0.012; cortex: r2 = 0.469, p= 0.01, respectively). Therefore, further downstream functional analysis focused on KIF1A and PLCB1 proteins.

Additional analysis confirmed the abundance of PLCB1 by Western blotting (Fig. 4f-h). The results showed that PLCB1 level decreased in the cortex (Fig. 4f) and hippocampus (Fig. 4g) in vehicle-treated CrT KO mice, while DCE-treated CrT KO mice showed a significant increase in PLCB1 levels in the cortex (p=0.0004) compared to the hippocampus and cerebellum (Fig. 4h).

DCE-mediated upregulation of PLCB1 levels in the cortex might lead to the production of inositol-1,4,5-triphosphate (IP3) (Rusciano *et al*, 2021), and IP3 modulates the NF-κβ pathway via dysregulation of a PKCα inhibitor, IκBα, thereby altering the abundance of NF-κβ-inducible genes. Thus, IκBα abundance was evaluated to determine whether the DCE-induced increase in PLCB1 levels affects targets downstream of this pathway. Western blotting indicated that IκBα (Fig. 4i) (p < 0.05) but not IκBβ (Fig. 4j) was significantly downregulated in the cortex in vehicle-treated CrT KO mice, while DCE treatment rescued IκBα protein levels in CrT KO mice (p=0.008), suggesting that PLCB1 involved in NF-κβ regulation in these mice.

## Discussion

The present study focuses on the CTD pathogenesis and the underlying mechanisms of DCE drug efficacy in CrT KO mouse model engineered to mimic the clinical features of CTD patients. Shotgun proteomics analysis, integrative bioinformatics and statistical modeling identified 14 key proteins that are dysregulated in the cerebellum, cortex, hippocampus and brain stem of *Slc6a8^-/v^* mice compared to WT mice and modulated by DCE. Notably, 13 of these proteins are related to ID disorders in human, including autism spectrum disorders (MECP2(Hammer *et al*, 2002; Shibayama *et al*, 2004; Wen *et al*, 2017), KIF1A (Dai *et al*, 2017; Esmaeeli Nieh *et al*, 2015; Lee *et al*, 2015; Ohba *et al*, 2015; Padiath, 2016; Tomaselli *et al*, 2017; Weller & Gartner, 2001), PLCB1 (Rusciano *et al*., 2021), NCAM1 (Arai *et al*, 2004; Atz *et al*, 2007; Gray *et al*, 2010; Varea *et al*, 2012), ANXA5 (Esmaeeli Nieh *et al*., 2015; Lee *et al*., 2015; Tomaselli *et al*., 2017)), bipolar disorder (FABP7 (Killoy *et al*, 2020), NCAM1, PLCB1 (Lo Vasco *et al*, 2013; Yang *et al*, 2016)), axonal neuropathy (DCLK1), ID (KIF1A (Ohba *et al*., 2015), L1CAM (Weller & Gartner, 2001)), regulation of neurite outgrowth (IGSF8 (Ray & Treloar, 2012)), leukodystrophy (LMNB1), cerebellar ataxia (LMNB1) (Dai *et al*., 2017; Padiath, 2016), abnormal behavior (PI4KA)(Pagnamenta *et al*, 2015), epileptic encephalopathies (PLCB1, MYO5A), or neurodegenerative diseases (ANK1) (De Jager *et al*, 2014).

Out of these 14 proteins, KIF1A abundance in the four brain regions (cortex, hippocampus, cerebellum and brain stem) was significantly correlated with DI in the object recognition test, while the abundance of this protein in the hippocampus, cortex and cerebellum correlated with the spontaneous alternation the Y-maze tests. Co-immunoprecipitation analysis using Western blotting confirmed that KIF1A interacts with PLCB1 in the various brain regions, suggesting a key role of KIF1A in CTD. The main observations of this study are summarized in Fig. 5. These results confirm what has been reported in the literature regarding the association of KIF1A and PLCB1 to cognitive function in different brain disorders (Lee *et al*., 2015; Manning *et al*, 2012)

**Fig. 5.**
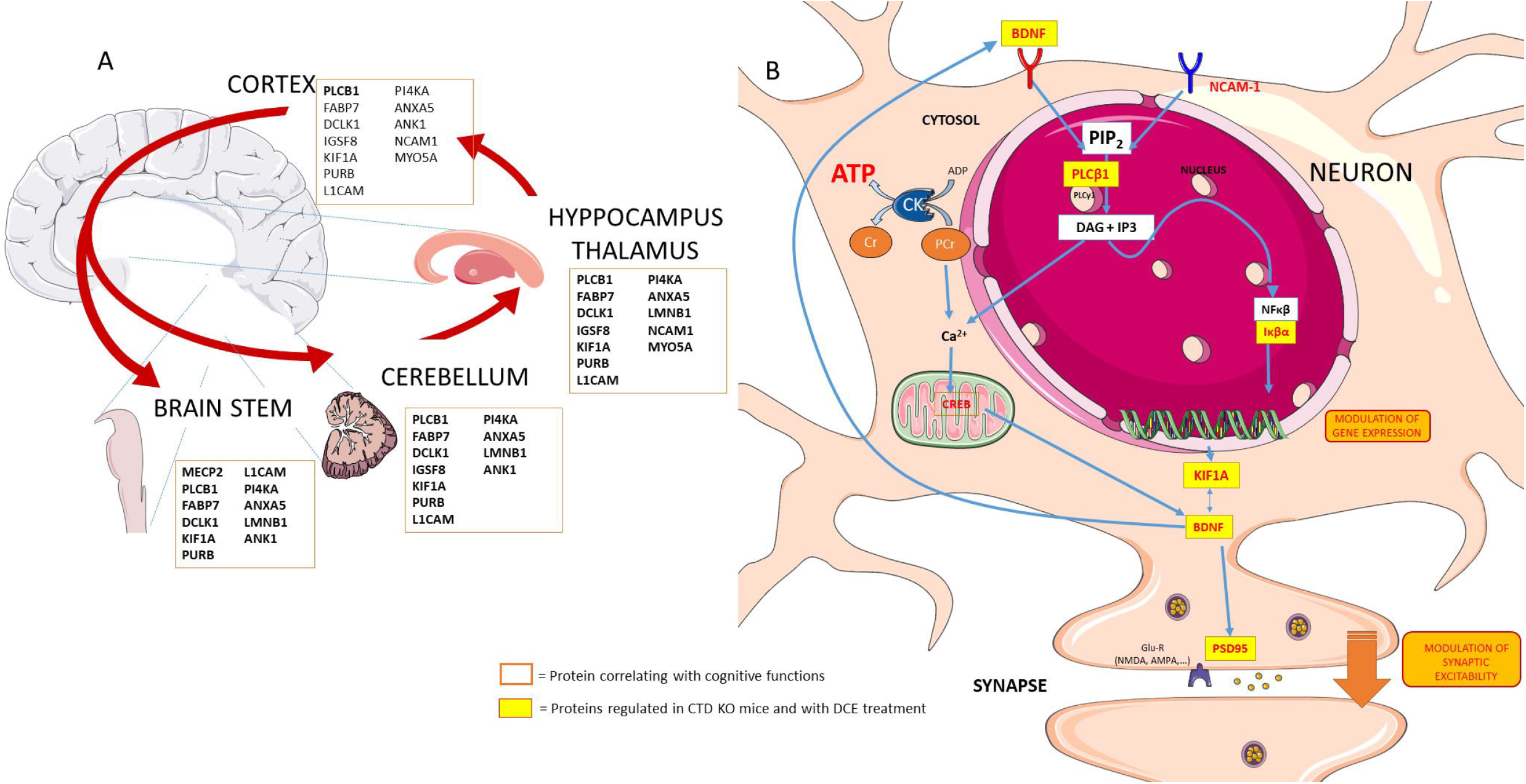
Schematic presentation showing the key players in the different brain regions and potential links between several proteins that are regulated in neurons in the context of CTD and by DCE treatment. **a,** Proteins regulated by DCE in CrT KO mice. **b**, DCE mediated the abundance of PLCB1. DCE-mediated upregulation of PLCB1 levels in the brain might lead to the production of inositol-1,4,5-triphosphate (IP3), and IP3 modulates the NF-κβ pathway via dysregulation of a PKCα inhibitor, IκBα, thereby altering the expression of NF-κβ-inducible genes which regulates the NF-κβ pathway by rescuing IκBα protein levels in CrT KO mice. NF-κβ is bound by IκBα and then translocates to the nucleus to activate target genes including KIF1A and BDNF.

KIF1A mutations have been found in patients with a severe neurodevelopmental disorder with Rett syndrome patients (Wang *et al*, 2019). KIF1A has been found to participate in vesicular transport. Vesicles containing the neurotrophin BDNF have been found to be under the control of KIF1A (Kondo *et al*., 2012). BDNF is produced in the neocortex throughout brain development and accelerates the overall redistribution of cortical neurons. DCE-mediated rescue of KIF1A levels in CrT KO mice resulted in higher levels of pro-BDNF/BDNF, which are linked to cognitive function improvement, suggesting that normal expression of KIF1A but not overexpression is indispensable for BDNF-mediated cognitive function.

We found that KIF1A interacts with PLCB1. PLCB1 plays a major role in vesicular trafficking within the cell, thus playing a direct role in axonal transport of neurotransmitters (Hannan *et al*, 2001). PLCB1 dysregulated signaling is linked to several brain disorders, including epilepsy, schizophrenia, bipolar disorder, Huntington’s disease, depression and Alzheimer’s disease (Alberini, 2009; Cholewa-Waclaw *et al*, 2016). We demonstrated that DCE modulates the relative abundance of PLCB1 in different brain regions. Our data showed that PLCB1 regulates the NF-κβ pathway by rescuing IκBα protein levels in CrT KO mice. Of note, NF-κβ signaling in the brain has been implicated in regulating neuronal survival and function (Kaltschmidt *et al*, 2006; Meffert *et al*, 2003). NF-κβ is bound by IκBα and then translocates to the nucleus to activate target genes. IκBα-deficient mice display deregulated and sustained NF-κβ activation (Lian *et al*, 2012), indicating a critical role for IκBα in NF-κβ regulation. CrT KO mice showed a decrease in IκBα abundance in the cortex and hippocampus, which was rescued by DCE treatment. This decrease in IκBα abundance probably regulates neuroinflammation as well as spatial memory formation and synaptic plasticity, probably through BDNF signaling. Our findings are in agreement with previous observations showing that stimulation of the neuroinflammatory response through NF-κβ activation may be therapeutically beneficial (Lian *et al*., 2012).

In conclusion, the results of the study indicated that the crosstalk between KIF1A and PLCB1 mediates cognitive function in the CTD. In addition, the results identified a panel of additional 12 proteins which suggests that Slc6a8-/y mice could have structural deficits including changes in neuronal migration or synaptogenesis. To our knowledge, this is the first study that describes some of the molecular mechanisms of CTD-related cognitive dysfunction and the therapeutic effect of DCE. Furthermore, the results of the study provide further evidence regarding the efficacy of DCE in treating the cognitive symptoms of CTD and restoring the abundance of key molecular players to normal levels in several brain regions. Little is known about the underlying mechanisms of the Cr-mediated behavioral deficits. Correlating proteomic changes to behavioral deficits provide mechanistic insights into Cr-mediated changes. Future studies can be designed to investigate this relationship. This study provides a shift in research paradigms and an advancement in intervention for CTD.

## Materials and methods

### Ethical considerations

All *in vivo* experiments were conducted in compliance with the European Communities Council Directive of 22 September 2010 and were approved by the Italian Ministry of Health (authorization number 259/2016-PR).

### Generation of CrT KO mice

Male CrT^-/v^ and CrT^+/v^ mice were generated on the C57BL/6J background as previously described (Baroncelli et *al*, 2014). The mice were housed at 22°C on a 12–12 h light–dark cycle and provided food and water *ad libitum*.

The presence of the *Slc6a8* mutation was confirmed by PCR as previously described (Baroncelli *et al*, 2016). Briefly, genomic DNA was isolated from tail tissue collected from P25 mice using the DNeasy^®^ Blood & Tissue kit from Qiagen according to the manufacturer’s protocol. The following primers were used for PCR amplification: F: AGGTTTCCTCAGGTTATAGAGA; R: CCCTAGGT GTATCTAACATCT; R1: TCGTGGTATCGTTATGCGCC. The amplicon sizes were as follows: CrT^+/v^ allele = 462 bp; mutant allele = 371 bp.

### DCE treatment

DCE was prepared as previously described (Trotier-Faurion *et al*., 2013). Ten milligrams of DCE was added to 0.375 g of Maisine^®^CC (Gattefossé) at room temperature, and then 125 mg of DHA (Sigma□Aldrich) was added. The mixture was vortexed for 5 min and shaken at 1000 × g in a thermomixer at 30°C for 48 h. Then, the sample was centrifuged at 20,000 × g for 10 min at room temperature, and the resulting supernatant was filtered through a 0.22 μm filter, placed in another tube and stored at +4°C prior to use.

DCE was intranasally administered to CrT KO mice for 30 days as previously reported (Ullio-Gamboa *et al*., 2019b), while wild-type (WT) and vehicle-treated mice were used as controls (N=8/group). 6 μL of DCE or vehicle (Maisine^®^CC with DHA) was placed in the nostril. The DCE (4 mg/g) or vehicle was given twice bilaterally (12 μL total volume).

### Behavioral testing

Behavioral testing started 14 days after the start of the treatment as was done previously (Baroncelli *et al*., 2016). Treatment continued during behavioral testing, which lasted two weeks, for a total of 30 days of treatment. Each mouse was subjected to all of the behavioral assessments in following order: the 24-h ORT (3 days), Y-maze test (1 day), and hidden platform MWM test (7 days).

#### ORT

The ORT apparatus consisted of a square arena (60 x 60 x 30 cm) made of polyvinyl chloride with black walls and a white floor as previously described (Baroncelli *et al*., 2016). The day before testing, the mice were individually habituated to the empty arena for 10 min. The ORT, which is based on the tendency of rodents to spend more time exploring a novel object than a familiar object, was used to measure short- and long-term memory and consisted of the sample phase and testing phase. During the sample phase, two identical objects were placed in diagonally opposite corners of the arena (approximately 6 cm from the walls), and the mice were allowed to explore the arena for 10 min. The testing phase was performed 24 h after the sample phase. An identical copy of one of the objects from the sample phase and a novel object were placed in the same locations, and the mice were returned to the arena and allowed to explore the objects for 5 min. The DI was calculated as follows: DI = (T new - T old)/(T new +T old), where T new is the time spent exploring the novel object and T old is the time spent exploring the familiar object (Baroncelli *et al*., 2016).

#### Y-maze spontaneous alternation test

The spontaneous alternation rate was measured using a Y-shaped maze with three symmetrical gray solid plastic arms at a 120-degree angle (26 cm long, 10 cm wide, and 15 cm high) as previously described (Baroncelli *et al*., 2016; Begenisic *et al*, 2014). The mice were placed in the center of the maze one at a time, and their movements were recorded for 8 min. The number of arm entries (all four limbs within an arm) and the number of triads (successive entries into all three arms) were recorded to calculate the spontaneous alternation percentage (defined as the number of triads divided by the number of possible alternations (total arm entries minus 2) multiplied by 100).

#### MWM test

The mice were subjected to 4 training trials per day for a total of 7 days. The apparatus consisted of a circular water tank (diameter, 120 cm; height, 40 cm) filled with water (23°C) to a depth of 25 cm. The water was made opaque by the addition of nontoxic white paint. Four starting positions arbitrarily designated the north (N), south (S), east (E), and west (W) positions were selected, thus dividing the tank into 4 quadrants. A square escape platform (11 × 11 cm) was submerged 0.5 cm below the water surface in the middle of one of the 4 quadrants. The mice were allowed to search for the escape platform for up to 60 s, and their swimming paths were automatically recorded by the Noldus Ethovision system. The last trial on the last training day was a probe trial, during which the escape platform was removed from the tank and the swimming paths of the mice were recorded for 60 s while they searched for the missing platform.

### Proteolysis and MS

Before MS analysis, total protein (15 μg) was extracted from tissues from the five tissues, i.e. muscle, cortical, cerebellar, hippocampal and brainstem tissues, then mixed with lithium dodecyl sulfate lysis buffer (Invitrogen), and incubated at 99°C for 5 min. Samples were then separated by electrophoresis for a short amount of time (5 min) at 200 V on a NuPAGE 4-12% Bis-Tris gel in 1X MES/SDS (Invitrogen) running buffer. The gels were stained with SimplyBlue SafeStain (Thermo) for 5 min followed by an overnight wash in water with gentle agitation. The band containing the whole proteome from each sample was excised from the polyacrylamide gel and treated as previously described (Hartmann & Armengaud, 2014). The proteins were in-gel proteolyzed with trypsin gold (Promega) in the presence of 0.01% Protease Max surfactant (Promega) at 50°C for 60 min. A total of 1 μL of the resulting peptide fraction (50 μL), corresponding to approximately 300 ng of peptide, was analyzed by liquid chromatography-tandem mass spectrometry (LC-MS/MS) using an Ultimate 3000 nano-LC system coupled to a Q-Exactive HF mass spectrometer (Thermo Scientific) as described previously (Hartmann & Armengaud, 2014). The peptides were loaded on a reverse-phase PepMap 100 C18 μ-precolumn (5 μm, 100 Å, 300 μm i.d. × 5 mm, Thermo Fisher) and then resolved on a nanoscale PepMap 100 C18 nanoLC column (3 μm, 100 Å, 75 μm i.d. × 50 cm, Thermo Fisher) at a flow rate of 0.2 μL.min^-1^ using a 90-min gradient (4% B from 0 to 3 min, 4-25% B from 3 to 78 min and 25-40% B from 78 to 93 min), with 0.1% HCOOH/100% H_2_O as mobile phase A and 0.1% HCOOH/80% CH3CN/20% H_2_O as mobile phase B. The mass spectrometer was operated in Top20 mode, with a scan range of 350 to 1800 *m/z*, and selection and fragmentation were performed using a 10 s dynamic exclusion time for the 20 most abundant precursor ions. Only ion precursors with a 2+ or 3+ charge were selected for HCD fragmentation, which was performed at a normalized collision energy of 27 eV.

### MS/MS spectra interpretation and differential proteomics

MS/MS spectra were assigned using Mascot Daemon software version 2.6.1 (Matrix Science) and the *Mus musculus* SwissProt database comprising 17,096 protein sequences. Peptide tolerance, MS/MS fragment tolerance, and the maximum number of missed cleavages were set to 5 ppm, 0.02 Da and 2, respectively. Carbamidomethylation of cysteine was considered a fixed modification, and oxidation of methionine was considered a variable modification. Peptides with p value ≤ 0.05 for homology threshold mode and proteins with at least two distinct peptides were selected (false discovery rate < 1%).

### Bioinformatics analysis of proteomics data

The general workflow of the bioinformatics analysis is shown in Supplemental Fig. 1. An in-house script was written using the R programming language to identify differentially expressed proteins between the three groups (WT, vehicle-treated CrT KO mice and DCE-treated CrT KO mice) in each of the five tissues, i.e. muscle, cortical, cerebellar, hippocampal and brainstem tissues. The following comparisons were analyzed: vehicle-treated CrT KO mice vs WT mice; DCE-treated CrT KO mice vs vehicle-treated CrT KO mice; and DCE-treated CrT KO mice vs WT. Initially, the proteomics data were normalized using variance stabilizing normalization (VSN) (Motakis *et al*, 2006). Proteins with at least 10 assigned MS/MS spectra across all samples were retained. An unsupervised variation filter was then applied to the proteomics data (Hamoudi *et al*, 2010), and samples of 8 proteins with MS/MS spectra were included. Differential abundance analysis of proteins among the different regions was carried out using a modified R package for reproducibility-optimized statistical testing (ROTS) (Suomi *et al*, 2017). The data were sorted according to the adjusted p value based on a false discovery rate < 0.05. Reproducibility plots and principal component analysis (PCA) were used to assess the quality of the separation of the data between the various groups that were being compared. The identified differentially expressed proteins were visualized using volcano plots and heatmaps. The heatmaps were generated using unsupervised hierarchical clustering carried out on the basis of Ward linkage and Euclidean distance to assess the degree of proteomic profile separation across the four brain regions among the three groups.

### Pathway analysis

Pathway analysis was performed to narrow down the differentially abundant proteins and identify their potential functions. To achieve this, the proteins whose abundances were significantly altered in CrT KO vehicle-treated mice compared with WT mice and in DCE-treated mice compared with vehicle-treated mice were selected for subsequent pathway analysis. The proteins whose abundances were altered in vehicle-treated mice and DCE-treated mice compared with WT mice were also selected. Pathway analysis using gene set enrichment was carried out using Enrichr (Chen *et al*, 2013; Kuleshov *et al*, 2016) focusing on the following sets: BioCarta_2016, Elsevier_Pathway_Collection, GO_Biological_Process_2018, GO_Molecular_Function_2018, KEGG_2019_Human, KEGG_2019_Mouse, MSigDB_Hallmark_2020, WikiPathways_2019_Mouse, WikiPathways_2019_Human, ClinVar_2019, DisGeNET, Jensen_DISEASES, and OMIM_Disease. Relevant pathways were selected based on a cutoff of p < 0.05.

### Statistical modeling of the differentially expressed proteomics data

**In order** to identify the patterns of differentially expressed proteins across the different regions, the data for the proteins identified from the pathway analysis were used to construct a multivariate statistical model using one-way ANOVA followed by Bonferroni’s post hoc test for comparisons among the cortex, cerebellum, brainstem, hippocampus and muscle. The proteins whose abundances were most markedly altered by the mutation and DCE treatment were selected. To identify the proteins involved in cognition, a stepwise regression model was constructed to assess the correlation between the levels of the differentially expressed proteins and performance in the ORT (DI) and the Y-maze test. The results were further validated using Pearson correlation analysis of the differentially abundant proteins among the different groups.

### Western blotting

Since PLCB1 and KIF1A abundance as well as their partners are associated to several brain disorders, Western blotting was used to determine their abundance as well as two inhibitors of NF-κB, IκBα and IκBβ, in dissected brain tissues. Briefly, brain tissues were homogenized in freshly prepared lysis buffer containing 20□mM Trizma-Base, 150□mM NaCl (pH 7.4) (Sigma□Aldrich, Saint-Quentin Fallavier, France), 1% Triton X-100, 4% complete protease inhibitor cocktail and 20% mix of anti-phosphatase inhibitors using a Precellys Evolution tissue homogenizer. The samples were then centrifuged at 2500 × g for l50min followed by 10000 x g for 20□min to obtain lysates for electrophoresis. The proteins (10 to 20 μg) and protein standards were mixed with Laemmli buffer and loaded on 4-15% Criterion TGX Stain-Free protein gels in 1×□TGS running buffer (all from Bio-Rad, Marnes-la-Coquette, France) and transferred to a 0.20μm PVDF membrane with the Trans-Blot Turbo RTA Midi Transfer Kit (Bio-Rad, Marnes-la-Coquette, France). The membranes were blocked for 30□min in 5% low-fat milk in TBS-0.1% Tween 20 at room temperature. The blots were probed with specific primary antibodies overnight at 4°C followed by horseradish peroxidase (HRP) secondary antibodies diluted 1:5000 or 1:50000 in 5% low-fat milk in TBS-0.1% Tween 20 at room temperature. For protein detection, the membranes were treated with ECL Prime Western Blotting reagent (Amersham, UK) or Clarity Western ECL Substrate and exposed with a ChemiDoc Touch Imaging System (Bio-Rad, Marnes-la-Coquette, France). The band density was quantified with Image Lab software (Bio-Rad, Marnes-la-Coquette, France). The following antibodies were used at the indicated dilutions: anti-PLCB1 (1:1000, Abeam, abl82359), anti-IκBα (1:500, Cell Signaling Technology, 4812S), anti-IκBβ (1:500, Cell Signaling Technology, 15519S), anti-PSD95 (1:2000, Merck, MABN68), anti-tubulin (1:2000, Sigma□Aldrich, T6199), and anti-KIF1A (1:1000, Abcam, ab180153).

### Co-IP

Protein extract samples (200 μg) from cortex and hippocampus were adjusted to a final volume of 600 μl with binding buffer (20 mM Tris-HCl (pH 7.5), 150 mM NaCl, 10% glycerol, 1 mM EDTA, 0.1% BSA, and 1X protease inhibitor) before the addition of 17.6 μl anti-KIF1A or anti-PLCB1 antibody and 100 U of Benzonase nuclease (Novagen 70746-3). The mixture was incubated overnight at 4°C on a rotating wheel. Forty-three microliters of Dynabeads protein G (Invitrogen, 10003D) were washed 3 times with PBS+0.05% Tween and once with binding buffer. The beads were then added to the immunoprecipitate and incubated for 1 h at room temperature with rotation. After incubation, the immunoprecipitate was washed twice with Benzonase buffer (20 mM Tris-HCl (pH 8.0), 20 mM NaC1, 10% glycerol, 2 mM MgCl2, 0.1% BSA, and 1X protease inhibitor (Roche)) and incubated in Benzonase buffer supplemented with 100 U of Benzonase nuclease for 30 min at 37°C before being washed three times with washing buffer (20 mM Tris-HCl (pH 7.5), 150 mM NaCl, 10% glycerol, 1 mM EDTA, 0.05% Tween, and 1X protease inhibitor). The immunoprecipitated proteins were eluted directly in 25 μl 1.5× Laemmli buffer supplemented with 200 mM DTT and 1 mM beta-mercaptoethanol at 95°C for 10 min before magnetic separation of the beads and MS. MS was carried out under similar conditions as those for the brain protein extracts except that the nano-UPLC gradient was reduced to 60 min.

## Supporting information

Supplemntal Figure 3 Table associated

## Data availability

The MS and proteomics dataset is available through the ProteomeXchange Consortium via the PRIDE partner repository (https://www.ebi.ac.uk/pride/) under dataset identifiers PXD024968 and 10.6019/PXD024968.

## Acknowledgments

This work was supported by Jerome Fondation Lejeune grant and by X-traordinaire, which is a patient group dedicated to rare intellectual disabilities.

## Author contributions

A.M. was responsible for project administration, conceptualization, funding acquisition and writing of the manuscript. F.C. and L.B. administered drugs to the mice, conducted behavioral studies and collected brain samples. E.P. administered drugs to the mice and performed genotyping and the behavioral studies. AC. and N.C. conducted biochemistry experiments. E.M. and F.C. conducted the Co-IP experiments. J.C.G. and J.A. designed and conducted the proteomic experiments. R.H. and R.A.H. conducted mathematical and statistical modeling of the proteomics data as well as bioinformatics analysis. M.R.S., R.H., L.B., J.A., R.A.H., H.B. and T.J. participated in the writing and review of the manuscript. All authors read and approved the final manuscript prior to submission.

## Additional information

**Mass spectrometry & proteomics data:** The reviewers may access this currently private dataset using https://www.ebi.ac.uk/pride/ website with the username as and password as 8bltDzWU.

---

**<u>Supplemental Fig. 1</u>: Flowchart depicting the workflow used to identify the key proteins that are associated with cognitive functions and regulated by DCE treatment in CTD pathogenesis.**

**1.** The raw data of 4035 proteins was generated using Q-exactive HF mass spectrometer. **2**. The raw proteomics data were normalized using the variance stabilizing normalization (VSN) function (Supplemental Table 1), and an unsupervised filter was applied. **3a.** The differentially abundant proteins between the three groups (WT, vehicle-treated CrT KO mice and DCE-treated CrT KO mice) in each brain region were identified using reproducibility-optimized statistical testing (ROTS) and sorting according to the adjusted p-value based on False Discovery Rate. 3b. The quality of the separation of the data between the various groups being compared was assessed using reproducibility plots and PCA. The differentially expressed proteins were visualized using volcano plots (Supplemental Fig. 2a-d). The degree of separation between the groups was assessed using unsupervised hierarchical clustering (Supplemental Fig. 3a-h). **4**. Venn diagram of the differentially expressed proteins was generated, and the overlapping proteins that showed a significant change in abundance in CrT KO mice compared with WT mice and in DCE-treated mice compared with vehicle-treated mice were selected for subsequent pathway analysis (Fig. 2a-d & Supplemental Table 2). Pathway analysis of the overlapping proteins was performed using Enrichr with a cutoff value of p < 0.05, and the proteins found to be involved in the different pathways and diseases were selected for further analysis. **5**. To identify the patterns of differentially expressed proteins across the different regions, the data of the proteins selected from the pathway analysis were used to construct a multivariate statistical model using one-way ANOVA followed by Bonferroni’s post hoc test for comparisons among the cortex, cerebellum, brainstem, hippocampus and muscle (Fig. 2e-f). **6**. The proteins whose abundances were most markedly altered by the mutation and DCE treatment were selected. Those most abundant proteins were significantly altered by the mutation compared to WT (vehicle-treated vs WT) and by the treatment compared to vehicle (DCE-treated vs. vehicle treated) and their abundance after treatment was restored to levels comparable to those in WT mice (Fig. 3a; Supplemental Tables 3 & 4). **7**. To identify the proteins involved in cognition, a stepwise regression model was constructed to assess the correlation between the levels of the differentially expressed proteins and performance in the ORT (DI) and the Y-maze test (Fig. 3b-c and Supplemental Table 5).

**<u>Supplemental Fig. 2</u>. Reproducibility plots, PCA and volcano plots for the four brain regions evaluated. a-d,** The differentially expressed proteins in the four brain regions (cortex (**a**), cerebellum (**b**), brainstem (**c**), and hippocampus (**d))** were visualized using volcano plots, and the quality of the separation of the data between the various groups being compared (vehicle-treated CrT KO mice vs. WT mice) and (DCE-treated vs. vehicle-treated CrT KO mice) was assessed using reproducibility plots and PCA.

**<u>Supplemental Fig. 3</u>. Unsupervised hierarchical clustering in the four brain regions.**

The degree of separation among the groups (vehicle-treated CrT KO mice, DCE-treated CrT KO mice, and WT mice) in the four brain regions (cortex (**a, b**), cerebellum (**c, d**), brainstem (**e, f**), and hippocampus (**g, h))** was assessed using unsupervised hierarchical clustering. The proteins shown on the clusters are listed in order in the table associated.

**<u>Supplemental Table 1</u>. List of detected proteins and their normalized abundances.**

**<u>Supplemental Table 2</u>. List of intersecting proteins ([DCE-treated vs. vehicle-treated] vs. [vehicle-treated vs. WT]) and ([DCE-treated vs. vehicle-treated] vs. [DCE vs. WT] vs. [vehicle-treated vs. WT]) extracted from the Venn diagram.**

**<u>Supplemental Table 3</u>. List and *p* values of the fourteen proteins that showed a significant change in abundance in the presence of the mutation and after treatment..**

A multivariate statistical model comparing WT mice with vehicle-treated CrT KO mice and DCE-treated CrT KO mice with vehicle-treated CrT KO mice was used to identify key proteins. Fourteen most abundant proteins were found to be significantly altered by the mutation compared to WT (vehicle-treated vs WT) and by the treatment compared to vehicle (DCE-treated vs. vehicle treated) and their abundance after treatment was restored to levels comparable to those in WT mice.

**<u>Supplemental Table 4</u>. Fold change in the abundance of the 14 proteins identified.**

**<u>Supplemental Table 5</u>. Proteins whose abundance was significantly correlated with the DI (a) and performance in the Y-maze test (b).**

## References

Alberini CM (2009) Transcription factors in long-term memory and synaptic plasticity. Physiol Rev 89: 121–145

Apostolo N, Smukowski SN, Vanderlinden J, Condomitti G, Rybakin V, Ten Bos J, Trobiani L, Portegies S, Vennekens KM, Gounko NV et al (2020) Synapse type-specific proteomic dissection identifies IgSF8 as a hippocampal CA3 microcircuit organizer. Nat Commun 11: 5171

Arai M, Itokawa M, Yamada K, Toyota T, Arai M, Haga S, Ujike H, Sora I, Ikeda K, Yoshikawa T (2004) Association of neural cell adhesion molecule 1 gene polymorphisms with bipolar affective disorder in Japanese individuals. Biol Psychiatry 55: 804–810

Atz ME, Rollins B, Vawter MP (2007) NCAM1 association study of bipolar disorder and schizophrenia: polymorphisms and alternatively spliced isoforms lead to similarities and differences. Psychiatr Genet 17: 55–67

Baroncelli L, Alessandri MG, Tola J, Putignano E, Migliore M, Amendola E, Gross C, Leuzzi V, Cioni G, Pizzorusso T (2014) A novel mouse model of creatine transporter deficiency. F1000Res 3: 228

Baroncelli L, Molinaro A, Cacciante F, Alessandri MG, Napoli D, Putignano E, Tola J, Leuzzi V, Cioni G, Pizzorusso T (2016) A mouse model for creatine transporter deficiency reveals early onset cognitive impairment and neuropathology associated with brain aging. Human Molecular Genetics 25: 4186–4200

Begenisic T, Baroncelli L, Sansevero G, Milanese M, Bonifacino T, Bonanno G, Cioni G, Maffei L, Sale A (2014) Fluoxetine in adulthood normalizes GABA release and rescues hippocampal synaptic plasticity and spatial memory in a mouse model of Down syndrome. Neurobiol Dis 63: 12–19

Braissant O, Henry H, Beard E, Uldry J (2011) Creatine deficiency syndromes and the importance of creatine synthesis in the brain. Amino Acids 40:1315–1324

Bruun TUJ, Sidky S, Bandeira AO, Debray FG, Ficicioglu C, Goldstein J, Joost K, Koeberl DD, Luisa D, Nassogne MC et al (2018) Treatment outcome of creatine transporter deficiency: international retrospective cohort study. Metab Brain Dis 33: 875–884

Cheillan D, Joncquel-Chevalier Curt M, Briand G, Salomons GS, Mention-Mulliez K, Dobbelaere D, Cuisset JM, Lion-Francois L, Portes VD, Chabli A et al (2012) Screening for primary creatine deficiencies in French patients with unexplained neurological symptoms. Orphanet J Rare Dis 7: 96

Chen EY, Tan CM, Kou Y, Duan Q, Wang Z, Meirelles GV, Clark NR, Ma’ayan A (2013) Enrichr: interactive and collaborative HTML5 gene list enrichment analysis tool. BMC Bioinformatics 14: 128

Cholewa-Waclaw J, Bird A, von Schimmelmann M, Schaefer A, Yu H, Song H, Madabhushi R, Tsai LH (2016) The Role of Epigenetic Mechanisms in the Regulation of Gene Expression in the Nervous System. J Neurosci 36: 11427–11434

Clark AJ, Rosenberg EH, Almeida LS, Wood TC, Jakobs C, Stevenson RE, Schwartz CE, Salomons GS (2006) X-linked creatine transporter (SLC6A8) mutations in about 1% of males with mental retardation of unknown etiology. Hum Genet 119: 604–610

Dai Y, Ma Y, Li S, Banerjee S, Liang S, Liu Q Yang Y, Peng B, Cui L, Jin L (2017) An LMNB1 Duplication Caused Adult-Onset Autosomal Dominant Leukodystrophy in Chinese Family: Clinical Manifestations, Neuroradiology and Genetic Diagnosis. Front Mol Neurosci 10: 215

De Jager PL, Srivastava G, Lunnon K, Burgess J, Schalkwyk LC, Yu L, Eaton ML, Keenan BT, Ernst J, McCabe C et al (2014) Alzheimer’s disease: early alterations in brain DNA methylation at ANK1, BIN1, RHBDF2 and other loci. Nat Neurosci 17:1156–1163

DesRoches CL, Patel J, Wang P, Minassian B, Salomons GS, Marshall CR, Mercimek-Mahmutoglu S (2015) Estimated carrier frequency of creatine transporter deficiency in females in the general population using functional characterization of novel missense variants in the SLC6A8 gene. Gene 565: 187–191

Esmaeeli Nieh S, Madou MR, Sirajuddin M, Fregeau B, McKnight D, Lexa K, Strober J, Spaeth C, Hallinan BE, Smaoui N et al (2015) De novo mutations in KIF1A cause progressive encephalopathy and brain atrophy. Ann Clin Transl Neurol 2: 623–635

Gray U, Dean B, Kronsbein HC, Robinson PJ, Scarr E (2010) Region and diagnosis-specific changes in synaptic proteins in schizophrenia and bipolar I disorder. Psychiatry Res 178: 374–380

Hachim MY, Hachim IY, Talaat IM, Yakout NM, Hamoudi R (2020) M1 Polarization Markers Are Upregulated in Basal-Like Breast Cancer Molecular Subtype and Associated With Favorable Patient Outcome. Front Immunol 11: 560074

Hammer S, Dorrani N, Dragich J, Kudo S, Schanen C (2002) The phenotypic consequences of MECP2 mutations extend beyond Rett syndrome. Ment Retard Dev Disabil Res Rev 8: 94–98

Hamoudi RA, Appert A, Ye H, Ruskone-Fourmestraux A, Streubel B, Chott A, Raderer M, Gong L, Wlodarska I, De Wolf-Peeters C et al (2010) Differential expression of NF-kappaB target genes in MALT lymphoma with and without chromosome translocation: insights into molecular mechanism. Leukemia 24: 1487–1497

Hannan AJ, Blakemore C, Katsnelson A, Vitalis T, Huber KM, Bear M, Roder J, Kim D, Shin HS, Kind PC (2001) PLC-beta1, activated via mGluRs, mediates activity-dependent differentiation in cerebral cortex. Nat Neurosci 4: 282–288

Hartmann EM, Armengaud J (2014) N-terminomics and proteogenomics, getting off to a good start. Proteomics 14: 2637–2646

Iki J, Inoue A, Bito H, Okabe S (2005) Bi-directional regulation of postsynaptic cortactin distribution by BDNF and NMDA receptor activity. Eur J Neurosci 22: 2985–2994

Jaggumantri S, Dunbar M, Edgar V, Mignone C, Newlove T, Elango R, Collet JP, Sargent M, Stockler-Ipsiroglu S, van Karnebeek CD (2015) Treatment of Creatine Transporter (SLC6A8) Deficiency With Oral S-Adenosyl Methionine as Adjunct to L-arginine, Glycine, and Creatine Supplements. Pediatr Neurol 53: 360–363 e362

Kaltschmidt B, Ndiaye D, Korte M, Pothion S, Arbibe L, Prullage M, Pfeiffer J, Lindecke A, Staiger V, Israel A et al (2006) NF-kappaB regulates spatial memory formation and synaptic plasticity through protein kinase A/CREB signaling. Mol Cell Biol 26: 2936–2946

Killoy KM, Harlan BA, Pehar M, Vargas MR (2020) FABP7 upregulation induces a neurotoxic phenotype in astrocytes. Glia 68: 2693–2704

Kondo M, Takei Y, Hirokawa N (2012) Motor protein KIF1A is essential for hippocampal synaptogenesis and learning enhancement in an enriched environment. Neuron 73: 7Q,3–7EΠ

Kuleshov MV, Jones MR, Rouillard AD, Fernandez NF, Duan Q, Wang Z, Koplev S, Jenkins SL, Jagodnik KM, Lachmann A et al (2016) Enrichr: a comprehensive gene set enrichment analysis web server 2016 update. Nucleic Acids Res 44: W90–97

Lee JR, Srour M, Kim D, Hamdan FF, Lim SH, Brunel-Guitton C, Decarie JC, Rossignol E, Mitchell GA, Schreiber A et al (2015) De novo mutations in the motor domain of KIF1A cause cognitive impairment, spastic paraparesis, axonal neuropathy, and cerebellar atrophy. Hum Mutat 36: 69–78

Lian H, Shim DJ, Gaddam SS, Rodriguez-Rivera J, Bitner BR, Pautler RG, Robertson CS, Zheng H (2012) IkappaBalpha deficiency in brain leads to elevated basal neuroinflammation and attenuated response following traumatic brain injury: implications for functional recovery. Mol Neurodegener 7: 47

Lo Vasco VR, Longo L, Polonia P (2013) Phosphoinositide-specific Phospholipase C ßl gene deletion in bipolar disorder affected patient. J Cell Commun Signal 7: 25–29

Lu B, Nagappan G, Lu Y (2014) BDNF and synaptic plasticity, cognitive function, and dysfunction. Handb Exp Pharmacol 220: 223–250

Manning EE, Ransome MI, Burrows EL, Hannan AJ (2012) Increased adult hippocampal neurogenesis and abnormal migration of adult-born granule neurons is associated with hippocampal-specific cognitive deficits in phospholipase C-beta1 knockout mice. Hippocampus 22: 309–319

Meffert MK, Chang JM, Wiltgen BJ, Fanselow MS, Baltimore D (2003) NF-kappa B functions in synaptic signaling and behavior. Nat Neurosci 6: 1072–1078

Motakis ES, Nason GP, Fryzlewicz P, Rutter GA (2006) Variance stabilization and normalization for one-color microarray data using a data-driven multiscale approach. Bioinformatics 22: 2547–2553

Ohba C, Haginoya K, Osaka H, Kubota K, Ishiyama A, Hiraide T, Komaki H, Sasaki M, Miyatake S, Nakashima M et al (2015) De novo KIF1A mutations cause intellectual deficit, cerebellar atrophy, lower limb spasticity and visual disturbance. J Hum Genet 60: 739–742

Padiath QS (2016) Lamin B1 mediated demyelination: Linking Lamins, Lipids and Leukodystrophies. Nucleus 7: 547–553

Pagnamenta AT, Howard MF, Wisniewski E, Popitsch N, Knight SJ, Keays DA, Quaghebeur G, Cox H, Cox P, Balla T et al (2015) Germline recessive mutations in PI4KA are associated with perisylvian polymicrogyria, cerebellar hypoplasia and arthrogryposis. Hum Mol Genet 24: 3732–3741

Raffaele M, Francesco C, Giulia S, Leonardo L, Mariangela G, Elena P, Grazia AM, Annarita F, Roberta B, Giovanni C et al (2020) Novel translational phenotypes and biomarkers for creatine transporter deficiency. Brain Commun

Ray A, Treloar HB (2012) IgSF8: a developmentally and functionally regulated cell adhesion molecule in olfactory sensory neuron axons and synapses. Mol Cell Neurosci 50: 238–249

Rusciano I, Marvi MV, Owusu Obeng E, Mongiorgi S, Ramazzotti G, Follo MY, Zoli M, Morandi L, Asioli S, Fabbri VP et al (2021) Location-dependent role of phospholipase C signaling in the brain: Physiology and pathology. Adv Biol Regul 79: 100771

Salomons GS, van Dooren SJ, Verhoeven NM, Cecil KM, Ball WS, Degrauw TJ, Jakobs C (2001) X-linked creatine-transporter gene (SLC6A8) defect: a new creatine-deficiency syndrome. Am J Hum Genet 68: 1497–1500

Shibayama A, Cook EH, Jr., Feng J, Glanzmann C, Yan J, Craddock N, Jones IR, Goldman D, Heston LL, Sommer SS (2004) MECP2 structural and 3’-UTR variants in schizophrenia, autism and other psychiatric diseases: a possible association with autism. Am J Med Genet B Neuropsychiatr Genet 128B: 50–53

Stockler S, Schutz PW, Salomons GS (2007) Cerebral creatine deficiency syndromes: clinical aspects, treatment and pathophysiology. Subcell Biochem 46:149–166

Suomi T, Seyednasrollah F, Jaakkola MK, Faux T, Elo LL (2017) ROTS: An R package for reproducibility-optimized statistical testing. Plos Computational Biology 13

Tomaselli PJ, Rossor AM, Horga A, Laura M, Blake JC, Houlden H, Reilly MM (2017) A de novo dominant mutation in KIF1A associated with axonal neuropathy, spasticity and autism spectrum disorder. J Peripher Nerv Syst 22: 460–463

Trotier-Faurion A, Dezard S, Taran F, Valayannopoulos V, de Lonlay P, Mabondzo A (2013) Synthesis and Biological Evaluation of New Creatine Fatty Esters Revealed Dodecyl Creatine Ester as a Promising Drug Candidate for the Treatment of the Creatine Transporter Deficiency. Journal of Medicinal Chemistry 56: 5173–5181

Trotier-Faurion A, Passirani C, Bejaud J, Dezard S, Valayannopoulos V, Taran F, de Lonlay P, Benoit JP, Mabondzo A (2015) Dodecyl creatine ester and lipid nanocapsule: a double strategy for the treatment of creatine transporter deficiency. Nanomedicine 10: 185–191

Ullio-Gamboa G, Udobi KC, Dezard S, Perna MK, Miles KN, Costa N, Taran F, Pruvost A, Benoit JP, Skelton MR et al (2019a) Dodecyl creatine ester-loaded nanoemulsion as a promising therapy for creatine transporter deficiency. Nanomedicine 14: 1579–1593

Ullio-Gamboa G, Udobi KC, Dezard S, Perna MK, Miles KN, Costa N, Taran F, Pruvost A, Benoit JP, Skelton MR et al (2019b) Dodecyl creatine ester-loaded nanoemulsion as a promising therapy for creatine transporter deficiency. Nanomedicine (Lond) 14: 1579–1593

Valayannopoulos V, Bakouh N, Mazzuca M, Nonnenmacher L, Hubert L, Makaci FL, Chabli A, Salomons GS, Mellot-Draznieks C, Brule E et al (2013) Functional and electrophysiological characterization of four non-truncating mutations responsible for creatine transporter (SLC6A8) deficiency syndrome. J Inherit Metab Dis 36: 103–112

van de Kamp JM, Mancini GM, Salomons GS (2014) X-linked creatine transporter deficiency: clinical aspects and pathophysiology. J Inherit Metab Dis 37: 715–733

Varea E, Guirado R, Gilabert-Juan J, Marti U, Castillo-Gomez E, Blasco-Ibanez JM, Crespo C, Nacher J (2012) Expression of PSA-NCAM and synaptic proteins in the amygdala of psychiatric disorder patients. J Psychiatr Res 46: 189–197

Wang J, Zhang Q, Chen Y, Yu S, Wu X, Bao X (2019) Rett and Rett-like syndrome: Expanding the genetic spectrum to KIF1A and GRIN1 gene. Mol Genet Genomic Med 7: e968

Weller S, Gartner J (2001) Genetic and clinical aspects of X-linked hydrocephalus (L1 disease): Mutations in the L1CAM gene. Hum Mutat 18: 1–12

Wen Z, Cheng TL, Li GZ, Sun SB, Yu SY, Zhang Y, Du YS, Qiu Z (2017) Identification of autism-related MECP2 mutations by whole-exome sequencing and functional validation. Mol Autism 8: 43

Yang YR, Kang DS, Lee C, Seok H, Follo MY, Cocco L, Suh PG (2016) Primary phospholipase C and brain disorders. Adv Biol Regul 61: 80–85

