## Supplementary figures and images for "Dodecyl Creatine Ester Improves Cognitive Function and Identifies Drivers of Creatine Deficiency"

### Supplemental Figure 1

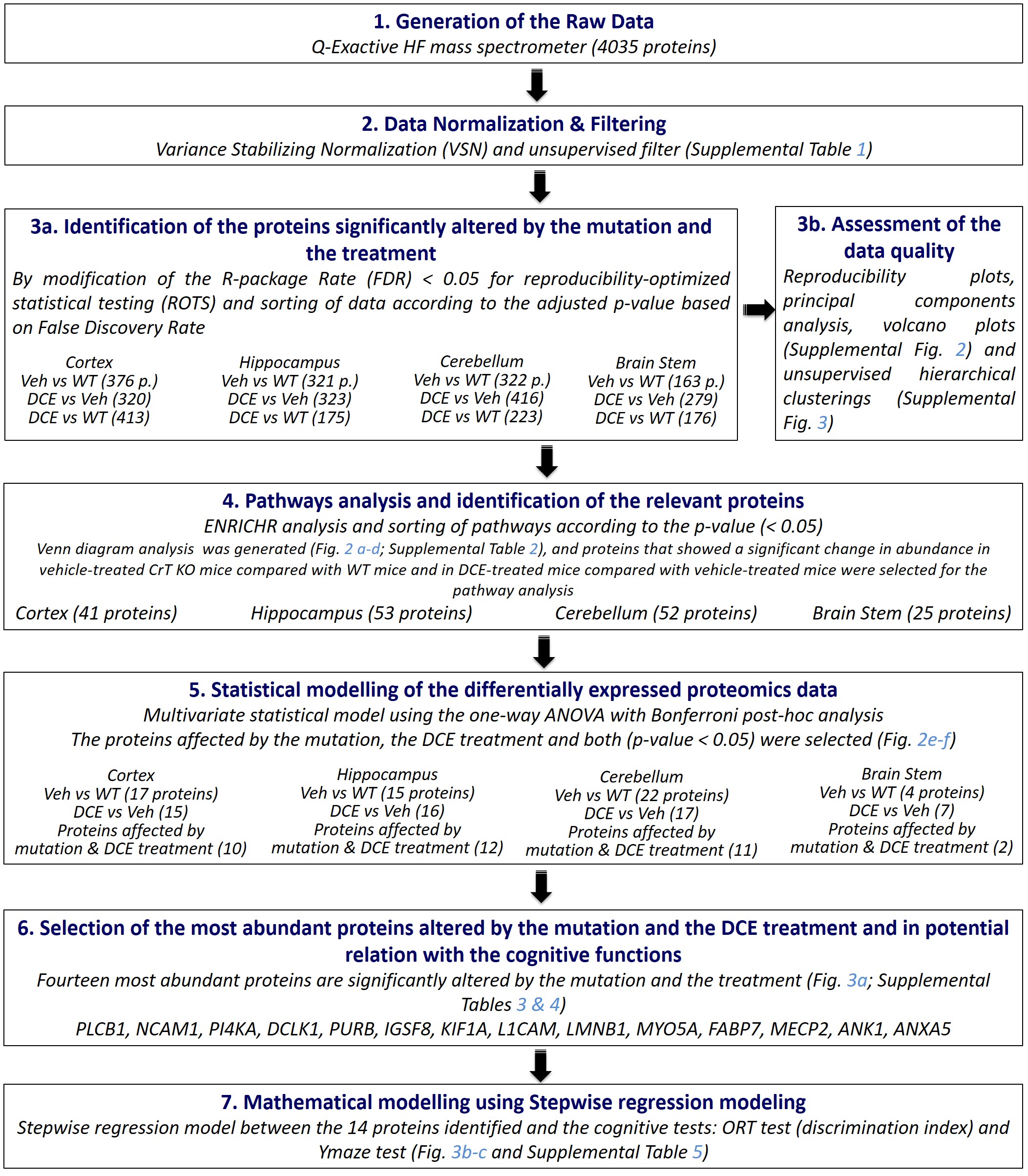

### Supplemental Figure 2

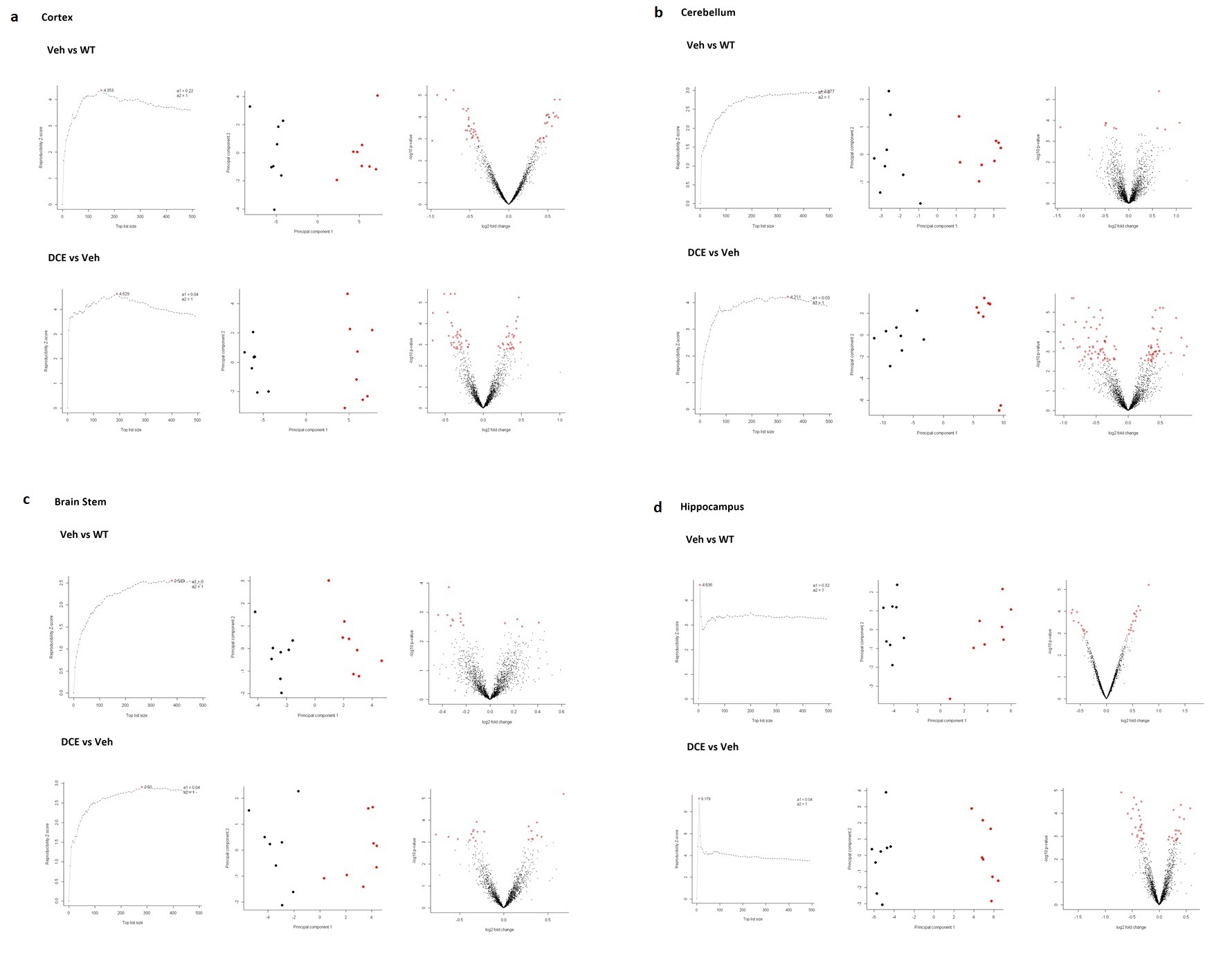

### Supplemental Figure 3a-d

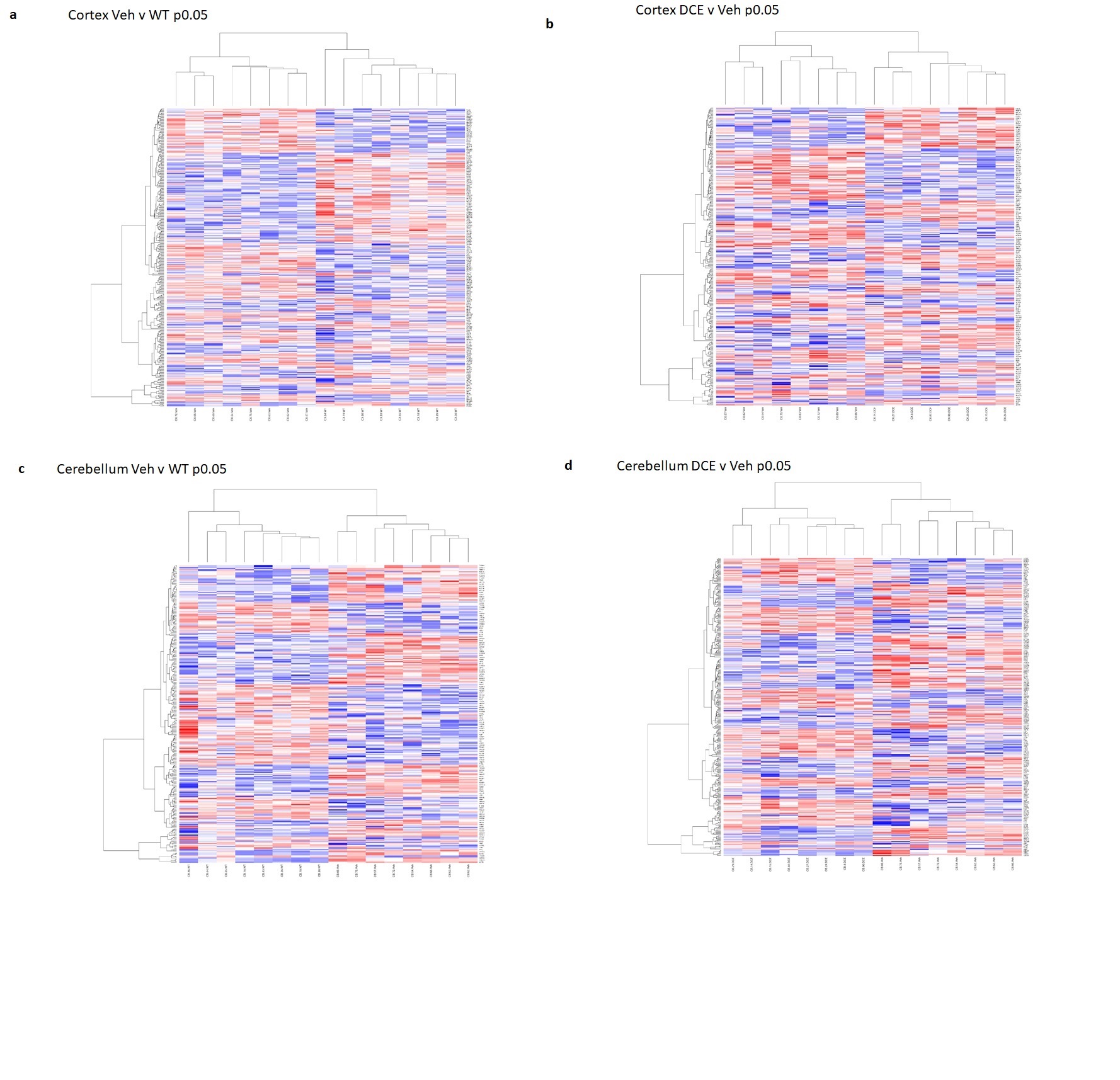

### Supplemental Figure 3e-h

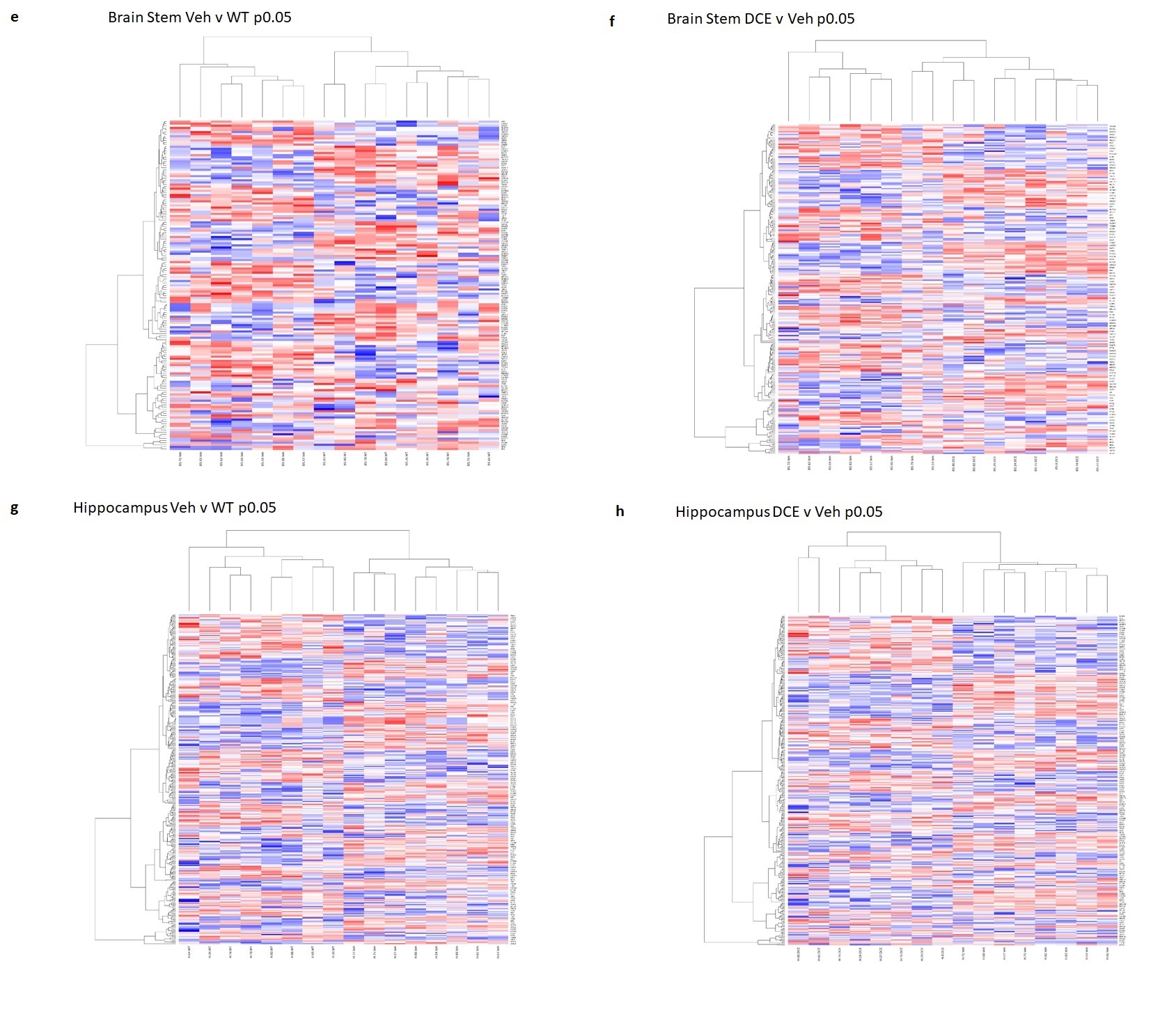
