## Supplementary material for "Dodecyl Creatine Ester Improves Cognitive Function and Identifies Drivers of Creatine Deficiency": Supplemntal Figure 3 Table associated

**Supplemental Figure 3: Table associated**

| **Lists of proteins shown in the unsupervised hierarchical clustering (Supplemental Figure 3)** | | | | | | | | | |
| --- | --- | --- | --- | --- | --- | --- | --- | --- | --- |
|  | **Cortex Veh v WT** | **Cortex**  **DCE v Veh** | **Hippocampus Veh v WT** | **Hippocampus DCE v Veh** | **Cerebellum Veh v WT** | **Cerebellum DCE v Veh** | **Cerebellum DCE v Veh** | **Brain Stem Veh v WT** | **Brain Stem DCE v Veh** |
| 1 | ECI2 | FRS1 | PDK | BCAT | PCDA | TICN | TICN | SR | GSK3 |
| 2 | AGO2 | FKBP8 | EFR3B | SET | CPSF7 | RENT1 | RENT1 | GRIA2 | DX39A |
| 3 | NOVA1 | KAD4 | THNS1 | NUDT3 | KNDC1 | ELMO1 | ELMO1 | DCUP | RAB5B |
| 4 | SAE1 | EXOC3 | SYCC | PLCL2 | RAE1L | KAD4 | KAD4 | NUDT4 | PA2G4 |
| 5 | HDGR3 | FPPS | PLCL2 | HOME1 | NUD10 | PPM1A | PPM1A | NUD10 | MARCS |
| 6 | DTNA | FKBP4 | KKCC2 | UBA5 | NUDT3 | PIMT | PIMT | PAPS1 | RELCH |
| 7 | COPG1 | FRIH | SNP47 | EFR3B | C170B | CE170 | CE170 | GORS2 | PGP |
| 8 | SF3A1 | LARP1 | INO1 | FKBP8 | PSB1 | NP1L4 | NP1L4 | IQEC3 | TOLIP |
| 9 | MYO1B | BRAF | CYLD | CSN2 | KBTB3 | VWA8 | VWA8 | RL23 | CAZA1 |
| 10 | HP1B3 | COPB2 | PSD11 | KPRB | GCST | ESYT1 | ESYT1 | TMX4 | GD1L1 |
| 11 | IMPCT | KIF3A | CACB2 | KKCC2 | K1C18 | BIG2 | BIG2 | RPN2 | PCKGM |
| 12 | CELF1 | CD82 | PGTA | EXC6B | K1C16 | APEH | APEH | OMGP | BLMH |
| 13 | NCLN | COX3 | STMN1 | F126B | K1C28 | INO1 | INO1 | XCT | MPP2 |
| 14 | NEB2 | VWA8 | A1AT5 | GGT7 | NEUL | PZP | PZP | CASC1 | PCY2 |
| 15 | GSTO1 | MPP3 | FKBP4 | PPR29 | TOM1 | CATD | CATD | OSBL9 | SRSF1 |
| 16 | LRC4B | UBA3 | PDK2 | NNRE | STK25 | KGP1 | KGP1 | RAE1 | RRAS2 |
| 17 | DDX42 | SNX5 | RO60 | VPS52 | G3BP1 | HCN2 | HCN2 | CNN3 | RS27 |
| 18 | HDGR2 | S2551 | CC50A | ASSY | GBG12 | CTNA1 | CTNA1 | ACYP1 | PUR8 |
| 19 | SI1L2 | HYES | CYB5B | AL9A1 | COQ9 | TMX2 | TMX2 | PRRT3 | SACS |
| 20 | RU17 | ANX11 | APOA1 | GNB5 | PCYXL | RS5 | RS5 | OXSR1 | STML2 |
| 21 | EIF3I | CX7A2 | CLIC4 | MYH1 | THEM6 | WDR37 | WDR37 | ACSA | CPT1A |
| 22 | ADK | FLOT1 | DDX42 | S14L2 | CDS1 | VAC14 | VAC14 | FGF1 | MUTA |
| 23 | STK26 | NACAM | CPT1A | RB11B | NIT1 | MK10 | MK10 | RENT1 | AL3B1 |
| 24 | ACSF2 | A1AT5 | P5CR2 | SYWC | SCRN3 | ACOT1 | ACOT1 | DOPD | ATPMD |
| 25 | MIF | RAP2C | GRM7 | NDUA8 | IKBL1 | RFOX1 | RFOX1 | PUF60 | ERMP1 |
| 26 | PGAM5 | SMU1 | PHOCN | NHRF1 | UBS3B | DPP10 | DPP10 | MECR | GPC5B |
| 27 | VCAM1 | AL1A3 | PGAM5 | ACO13 | LYPA2 | CBPE | CBPE | XXLT1 | SYNE1 |
| 28 | CBS | SRSF6 | CNR1 | EFHD2 | ASAH1 | NAC1 | NAC1 | TOM34 | HNRPF |
| 29 | KIF7 | RU17 | KIT | DGKB | YTHD2 | NU4M | NU4M | AL3A1 | COTL1 |
| 30 | PLCL1 | PSB5 | LYAG | ATPMD | QCR6 | S12A4 | S12A4 | CPT1A | PZP |
| 31 | ENDD1 | IRGQ | WDFY3 | RS25 | ECI1 | SCN1A | SCN1A | PEPD | MLP3A |
| 32 | PI3R4 | KCTD8 | A1AT3 | CAMP3 | ARL3 | NDUA8 | NDUA8 | ENTP2 | DCUP |
| 33 | CX7A2 | TRI62 | A1AT2 | ACAD8 | IPO9 | TNPO1 | TNPO1 | CUL2 | IDI1 |
| 34 | NDUB6 | SYSM | NNRE | SRGP2 | NTRI | G37L1 | G37L1 | SHRM2 | NTRI |
| 35 | HYES | EPHA6 | GRHPR | RRAS2 | RS9 | ICMT | ICMT | SV2B | TIM29 |
| 36 | SCRN3 | PPM1B | PHAR1 | OST48 | GPSM1 | ITSN1 | ITSN1 | NDUB4 | FKBP3 |
| 37 | KAD4 | TOM1 | NAA15 | CTND1 | 6PGL | ABCB7 | ABCB7 | NDUS5 | TOM40 |
| 38 | ITPR3 | EIF3F | NIPS2 | SF3B3 | RUVB1 | RIDA | RIDA | RL21 | GLYM |
| 39 | F126B | NCLN | PZP | L2GL1 | ACOT1 | ACO13 | ACO13 | RRAS2 | HMOX2 |
| 40 | DIAP1 | EF1B | NGEF | OPLA | EPMIP | AP1G1 | AP1G1 | MIRO2 | K1C15 |
| 41 | FPPS | CKAP4 | PSMD5 | FERM2 | PKN1 | BORG4 | BORG4 | GLYM | K2C74 |
| 42 | PADI2 | RS27 | KAD4 | GCN1 | PURG | GBRA6 | GBRA6 | GD1L1 | K2C1 |
| 43 | RO60 | FSD1 | RRAS2 | KIF1C | NDUB4 | GRD2I | GRD2I | RASH | T120A |
| 44 | SNX6 | COR2B | THTM | ITB1 | BCR | MLC1 | MLC1 | NAC2 | CHMP3 |
| 45 | ZNT9 | VCAM1 | CBS | TEN2 | PPM1B | AL9A1 | AL9A1 | PITM1 | RAE1 |
| 46 | NDUB7 | SRSF4 | C170B | RL8 | RBM3 | ANXA7 | ANXA7 | LAT1 | CTRO |
| 47 | CAMP2 | CBS | CAN5 | TXTP | PPIB | KPCA | KPCA | LIPA2 | PRRT3 |
| 48 | VP13A | RRAS2 | H2AY | NOMO1 | SYLC | CLIP1 | CLIP1 | ITIH3 | ABCB9 |
| 49 | INF2 | MYO1B | OX2G | MYEF2 | RBM39 | SYIC | SYIC | GPC5B | PRVA |
| 50 | ANX11 | TENA | NEUL | KCC4 | XPP1 | ACTN2 | ACTN2 | DIP2A | PUF60 |
| 51 | MTX1 | BPHL | K1C14 | ATAD3 | RFOX1 | GNA13 | GNA13 | SPART | GBRL2 |
| 52 | CSN2 | HEXB | K1C13 | AMPD2 | NDRG3 | MK09 | MK09 | A1AT5 | DTNA |
| 53 | SGT1 | PIMT | K1C15 | PGTA | VPS29 | IPYR | IPYR | A1AT | H10 |
| 54 | FAK1 | SPN90 | MARE1 | F1712 | TMX2 | OTUB1 | OTUB1 | STK39 | MECR |
| 55 | SRGP3 | QCR8 | CKAP4 | PDE4D | WDR37 | TMOD2 | TMOD2 | KBTBB | PP2BB |
| 56 | A1AT5 | SGT1 | NDRG3 | SYT7 | PCYOX | NDUAD | NDUAD | CHRD1 | HPRT |
| 57 | NLRC5 | STXB5 | RHG39 | S2551 | CBPE | S12A6 | S12A6 | PLIN3 | ARP3 |
| 58 | PRP19 | RINI | FABP7 | KPRA | BAG6 | ABLM2 | ABLM2 | PCS1N | UBP14 |
| 59 | MCCB | MTMR1 | SRBS1 | UBP15 | RENT1 | NDUS7 | NDUS7 | FAHD2 | CAP1 |
| 60 | A1AT3 | SRGP3 | DHX15 | RAB18 | S100B | CX6B1 | CX6B1 | UB2D2 | VAT1L |
| 61 | A1AT2 | COX6C | SF3B3 | TMX3 | PLXB1 | PSA4 | PSA4 | SGTA | DDX5 |
| 62 | FERM2 | YES | ALG2 | FCSK | NP1L1 | SYRC | SYRC | DHB11 | COX41 |
| 63 | AOFB | VIME | SE6L1 | NAKD2 | FAHD2 | FABP7 | FABP7 | HACL2 | SFXN3 |
| 64 | RAP1B | PIN1 | LRRC7 | RO60 | KAD4 | DDAH2 | DDAH2 | DHRS1 | KIF5A |
| 65 | TSN2 | SYFA | KIF1B | CYTC | A1AT | BASP1 | BASP1 | RRBP1 | LONM |
| 66 | SYNPR | PPIB | NUDC | MBOA7 | F162A | MK14 | MK14 | SPA3K | GNAI2 |
| 67 | ELMO2 | NOMO1 | AMPB | AL1A2 | ABCB7 | MK12 | MK12 | CRYAB | NDUS3 |
| 68 | NDUV2 | ASAP2 | AHSA1 | RBM3 | SPRE | PSB6 | PSB6 | RAB5B | PAK1 |
| 69 | NDUBB | KCD16 | VPS51 | AGRB1 | AL9A1 | SCN5A | SCN5A | GNA14 | PURB |
| 70 | SYK | SFXN5 | SYWC | ROCK1 | KCNC3 | C170B | C170B | PRS4 | RS4X |
| 71 | PPID | THEM4 | HARS1 | KC1D | THEM4 | NDUAB | NDUAB | SFXN1 | COR1A |
| 72 | SPN90 | TXNL1 | RB11A | DIP2B | NNRE | MUC18 | MUC18 | IMPCT | HNRPL |
| 73 | CAN2 | 6PGD | SPN90 | VP13A | BPHL | THIKB | THIKB | JIP4 | NCAM2 |
| 74 | GSTM3 | PPM1A | NCEH1 | RSMB | IPYR | NUDT4 | NUDT4 | PSA7 | DESM |
| 75 | NDUA4 | GFAP | EXOC8 | ACACA | A1AT1 | RAE1L | RAE1L | MP2K2 | CSN4 |
| 76 | XPO1 | SNG3 | GNB5 | CKAP4 | A1AT4 | TOM1 | TOM1 | NOMO1 | CNTP2 |
| 77 | CLD11 | AHSA1 | PLCB1 | CAH2 | HCDH | RL19 | RL19 | FABP7 | AT2A1 |
| 78 | RL13A | PLST | DNJA1 | PLST | HPLN1 | G3BP1 | G3BP1 | GNA11 | SYDC |
| 79 | ARL8B | PPID | CDC37 | CUL3 | NDRG1 | PLCE | PLCE | PPCE | MYH11 |
| 80 | CCG8 | TTC7B | RPGP1 | PYGL | APOA1 | CTND1 | CTND1 | RS18 | FABP5 |
| 81 | CYBP | RPGP2 | RPGP2 | VAMP1 | ILEUA | NDUB9 | NDUB9 | LIPA3 | PTPA |
| 82 | PRRT3 | KCAB2 | PLXA2 | CSN4 | PZP | PDE1B | PDE1B | PA1B2 | RAB6B |
| 83 | PDIA4 | GSLG1 | CCAR2 | SYYC | TRAP1 | NDUB7 | NDUB7 | THOP1 | KAPCB |
| 84 | SAM50 | DGKB | C1TM | PI42A | AL1A7 | NDUB4 | NDUB4 | ACOT9 | PRS6B |
| 85 | K1107 | SCN2A | M4K4 | SNAA | OAT | S38A3 | S38A3 | LSAMP | K2C73 |
| 86 | ASTN1 | CX6B1 | ELMO2 | CAP2 | USO1 | NAMPT | NAMPT | SC6A1 | GBB4 |
| 87 | SLK | MARCS | ACADV | IGSF8 | LPPRC | ARRD1 | ARRD1 | NDUAC | NECP1 |
| 88 | SYTC | ACOC | NCHL1 | KCY | TRIM2 | IKBL1 | IKBL1 | A1AT1 | HNRDL |
| 89 | ATIF1 | PPCE | VAT1 | DDX3L | AFG32 | BROX | BROX | A1AT3 | PSA7 |
| 90 | CE170 | ACTN2 | DOCK4 | HNRPQ | DHPR | OPA3 | OPA3 | A1AT4 | ENPP6 |
| 91 | STRN3 | ACAD9 | S12A2 | H31 | AUHM | UBXN6 | UBXN6 | A1AT2 | DP13A |
| 92 | SAE2 | KCY | DGKG | DEST | NIPS2 | SRGP3 | SRGP3 | CO3 | AT2A3 |
| 93 | FYN | ACBG1 | SYIM | PSMD2 | PDIA4 | ECI1 | ECI1 | UBQL2 | FAAA |
| 94 | DMWD | PP1R7 | PRPS1 | EIF3A | CDC37 | CDS1 | CDS1 | TRXR1 | GSTM3 |
| 95 | AL4A1 | DCTN2 | LNEBL | LIPA2 | FKBP4 | CNN3 | CNN3 | MAOM | NDUA4 |
| 96 | LRC47 | SEP4 | AFG32 | DDX3Y | FLNA | RMD3 | RMD3 | CDC37 | SNPH |
| 97 | ARBK1 | RAB7A | PI42B | NT5D3 | SYUA | DGKG | DGKG | GSTM3 | A4 |
| 98 | TXTP | SYIM | VISL1 | TCPW | PRS6B | SCRN3 | SCRN3 | FERM2 | PGCA |
| 99 | RLA0 | INP4A | PDXK | TPM1 | S27A1 | RS9 | RS9 | AL3A2 | CD9 |
| 100 | NCHL1 | RL15 | PI4KA | CADM4 | TFR1 | SNRPA | SNRPA | FKBP4 | VPP1 |
| 101 | NOMO1 | ARFG1 | CAP2 | CSRP1 | SNX27 | NEK6 | NEK6 | HPLN4 | PYGL |
| 102 | MARE1 | CTTB2 | IGSF8 | OLA1 | ALG2 | TPM2 | TPM2 | FLNA | AT5F1 |
| 103 | PIN1 | GDAP1 | VIME | BAIP2 | BASP1 | CPSF5 | CPSF5 | PZP | KINH |
| 104 | AL1A1 | RSSA | PCBP1 | PCCB | PPCE | MIRO2 | MIRO2 | PGCA | PDXK |
| 105 | AT8A1 | AL1B1 | LMNB1 | TPM2 | FABP7 | NDUS8 | NDUS8 | ILEUA | VDAC3 |
| 106 | H2A1B | SYLC | PRDX1 | UCRI | MARCS | MARE1 | MARE1 | PP2AA | COF1 |
| 107 | LONM | PLXA1 | PAK3 | RL18 | GBB2 | KCTD8 | KCTD8 | PP2AB | SNAB |
| 108 | NONO | DLG1 | PAK1 | TRIM3 | VATH | PURG | PURG | MP2K1 | GLSK |
| 109 | KAPCA | RAB14 | NAC1 | CCAR2 | IF4A1 | TOM22 | TOM22 | NCAM2 | SPTB1 |
| 110 | THOP1 | CSKP | LIPA2 | NRX2A | VATE1 | CNTN2 | CNTN2 | DPP6 | TLN1 |
| 111 | PTPRZ | KPCE | L1CAM | PPCE | LRP1 | S6A17 | S6A17 | VISL1 | PFKAP |
| 112 | DDX5 | ROCK2 | MINK1 | ACAD9 | PFKAL | KCND2 | KCND2 | HPRT | SYNJ1 |
| 113 | HNRPM | PSMD2 | PPM1E | GELS | KINH | LNEBL | LNEBL | GNAI3 | BCAS1 |
| 114 | CEND | ARHG2 | FUBP2 | PLXA2 | ANXA5 | H10 | H10 | VATC1 | NFL |
| 115 | SGIP1 | ARP2 | TNIK | COPA | AMPH | OPLA | OPLA | KAP3 | NFM |
| 116 | NCAM2 | RS3 | KCY | WFS1 | SEP8 | COTL1 | COTL1 | SFXN5 | NFH |
| 117 | DREB | IGSF8 | BCAS1 | SUCB1 | OXR1 | FKBP4 | FKBP4 | BASP1 | MPCP |
| 118 | H2AX | ADA22 | PPCE | AMPH | TAGL3 | SYIM | SYIM | PPCEL | VATB2 |
| 119 | AK1A1 | DYL2 | SLK | TCPD | PEBP1 | OAT | OAT | KCY | KCRU |
| 120 | PSD3 | HYOU1 | GSTM5 | SUCA | SFXN5 | CAP1 | CAP1 | COF2 |  |
| 121 | SYGP1 | KI21A | NT5D3 | H4 | ARC1A | ACBP | ACBP | TMOD2 |  |
| 122 | TAGL3 | PARK7 | NEB2 | SYAC | SODM | DHPR | DHPR | CTNA2 |  |
| 123 | CUL3 | CAH2 | CSRP1 | PTPRZ | VATC1 | TRAP1 | TRAP1 | CADM2 |  |
| 124 | KI21A | CLIP2 | UGPA | KIF1A | KIF1A | IPO5 | IPO5 | ETFD |  |
| 125 | MAG | CPNE6 | CADM3 | TCPZ | PURB | USO1 | USO1 | USO1 |  |
| 126 | ALDH2 | TCPB | XPO2 | PRDX6 | CAH2 | FLNA | FLNA | IPO5 |  |
| 127 | MOG | STIP1 | CADM4 | MP2K1 | VATB2 | NSF1C | NSF1C | AL1A7 |  |
| 128 | NFH | PFKAL | SAC1 | SNP25 | NCAM1 | CTND2 | CTND2 | CALB2 |  |
| 129 | ROA2 | PYGM | KIF1A | ACLY | SCOT1 | ARG33 | ARG33 | MYH11 |  |
| 130 | NRCAM | ACLY | LRP1 | HNRH2 | DPYL1 | SHAN1 | SHAN1 | PABP1 |  |
| 131 | MATR3 | NRCAM | FSCN1 | DDX3X | PYGM | GNAS1 | GNAS1 | COR1C |  |
| 132 | GRIN1 | SYAC | TLN2 | PURB | PYGB | PPCE | PPCE | LRP1 |  |
| 133 | PABP1 | ECHA | NRCAM | CALM1 | 2AAA | ACADV | ACADV | PARK7 |  |
| 134 | IDHC | ACTN4 | MY18A | RAC1 | VIME | SAM50 | SAM50 | SCPDL |  |
| 135 | ARF3 | RAB10 | PDIA3 | BDH | HS74L | CTNA2 | CTNA2 | CAN2 |  |
| 136 | PYGM | BDH | SYT2 | IDHC | HBB1 | SC6A1 | SC6A1 | VDAC3 |  |
| 137 | SIR2 | SIR2 | ENPL | GRIN1 | THIL | KIF1A | KIF1A | NDUS3 |  |
| 138 | GPM6B | 6-Sep | PYGM | L1CAM | DPYL5 | FUS | FUS | PHB2 |  |
| 139 | PLCB1 | TPPP | DPP6 | TLN1 | ACTS | GDIR1 | GDIR1 | PRDX5 |  |
| 140 | CELF2 | PLCB1 | QCR2 | NAC1 | HS90A | CNDP2 | CNDP2 | MYH14 |  |
| 141 | ILF3 | SH3G2 | HS105 | TENA | ALDOA | KCY | KCY | EAA1 |  |
| 142 | ATP5I | NCKP1 | SPTB1 | NRCAM | TBB2B | NDUS3 | NDUS3 | NDUS2 |  |
| 143 | SYLC | NFASC | PDE2A | MY18A | ACTB | PHB2 | PHB2 | AL1A1 |  |
| 144 | CBPE | ACTN1 | TCPA | FSCN1 |  | SEP5 | SEP5 | MYO1D |  |
| 145 | DNPEP | KAP3 | BIN1 | HS105 |  | TCPA | TCPA | DHPR |  |
| 146 | MARCS | FAS | RP3A | SH3G2 |  | SEP4 | SEP4 | CAH2 |  |
| 147 | STRN | PACN1 | TPP2 | SYT2 |  | PRDX6 | PRDX6 | ANXA5 |  |
| 148 | AL1B1 | TENR | PGM2L | SRCN1 |  | ANXA5 | ANXA5 | HS105 |  |
| 149 | PCCB | CAPS1 | SIR2 | NFASC |  | LONM | LONM | DPYL4 |  |
| 150 | KCAB2 | SCOT1 | FAK2 | E41L3 |  | GPDA | GPDA | GPDM |  |
| 151 | RPN1 | HS90A | SCOT1 | AATC |  | BDH | BDH | GNAO |  |
| 152 | CSRP1 | MAP1A | DPYL5 | DLDH |  | VDAC2 | VDAC2 | SCOT1 |  |
| 153 | ATP5L | ACTS | CNTN1 | DPYL4 |  | IF4A1 | IF4A1 | NFH |  |
| 154 | KCRS | DPYL3 | MYO5A | AATM |  | IDHC | IDHC | PYGM |  |
| 155 | AP3M2 | NCAM1 | PYGB | PP2BA |  | VDAC3 | VDAC3 | EAA2 |  |
| 156 | TPM3 | VATA | KCRU | TENR |  | GFAP | GFAP | NFM |  |
| 157 | PPCE | AATM | HS90B | DPYL1 |  | NDUS2 | NDUS2 | NFL |  |
| 158 | WDR37 | ALBU | ENOG | MAP1B |  | PYGL | PYGL |  |  |
| 159 | SYNPO | G3P | DPYL3 | HS90A |  | RAN | RAN |  |  |
| 160 | LMNB1 | ACTB | DYHC1 | ENOG |  | ARC1A | ARC1A |  |  |
| 161 | ACTN3 |  |  | ENOA |  | MECP2 | MECP2 |  |  |
| 162 | BAIP2 |  |  | DYHC1 |  | KIF5A | KIF5A |  |  |
| 163 | AMPL |  |  |  |  | INP4A | INP4A |  |  |
| 164 | PPME1 |  |  |  |  | CKAP5 | CKAP5 |  |  |
| 165 | HPCL4 |  |  |  |  | TLN1 | TLN1 |  |  |
| 166 | ANXA5 |  |  |  |  | GPDM | GPDM |  |  |
| 167 | PI42A |  |  |  |  | SAHH3 | SAHH3 |  |  |
| 168 | TSN7 |  |  |  |  | ACSL6 | ACSL6 |  |  |
| 169 | PI4KA |  |  |  |  | 1433B | 1433B |  |  |
| 170 | COX2 |  |  |  |  | SH3G2 | SH3G2 |  |  |
| 171 | THIL |  |  |  |  | KIF5C | KIF5C |  |  |
| 172 | NEUM |  |  |  |  | HBA | HBA |  |  |
| 173 | LIPA3 |  |  |  |  | H13 | H13 |  |  |
| 174 | ACTN4 |  |  |  |  | H14 | H14 |  |  |
| 175 | MBP |  |  |  |  | NPTN | NPTN |  |  |
| 176 | PLEC |  |  |  |  | QCR2 | QCR2 |  |  |
| 177 | HS105 |  |  |  |  | MIC60 | MIC60 |  |  |
| 178 | ENOB |  |  |  |  | NDUS1 | NDUS1 |  |  |
| 179 | NFL |  |  |  |  | MYH9 | MYH9 |  |  |
| 180 | NFM |  |  |  |  | CNTN1 | CNTN1 |  |  |
| 181 | SCOT1 |  |  |  |  | S12A5 | S12A5 |  |  |
| 182 | TAU |  |  |  |  | DPYL4 | DPYL4 |  |  |
| 183 | DCLK1 |  |  |  |  | VATB2 | VATB2 |  |  |
| 184 | DPYL3 |  |  |  |  | VDAC1 | VDAC1 |  |  |
| 185 | NCAM1 |  |  |  |  | DPYL1 | DPYL1 |  |  |
| 186 | ALBU |  |  |  |  | PFKAM | PFKAM |  |  |
| 187 | HS90A |  |  |  |  | ANK1 | ANK1 |  |  |
| 188 | ENOG |  |  |  |  | PYGM | PYGM |  |  |
| 189 |  |  |  |  |  | ACTS | ACTS |  |  |
| 190 |  |  |  |  |  | AT2B2 | AT2B2 |  |  |
| 191 |  |  |  |  |  | AINX | AINX |  |  |
| 192 |  |  |  |  |  | TBB2B | TBB2B |  |  |
| 193 |  |  |  |  |  | TBB5 | TBB5 |  |  |
| 194 |  |  |  |  |  | TBA1C | TBA1C |  |  |
| 195 |  |  |  |  |  | TBA1A | TBA1A |  |  |
