## Supplemental Table 3 for "Dodecyl Creatine Ester Improves Cognitive Function and Identifies Drivers of Creatine Deficiency"

**Supplemental Table 3: List and *p* values of the fourteen proteins that showed a significant change in abundance in the presence of the mutation and after treatment.**

**Supplemental Table 3a**

| **Brain Region** | **Protein** | **Group Comparison** | **N number** | **Statistics** | **p value / Significance** |
| --- | --- | --- | --- | --- | --- |
| **Cortex** | PLCB1^31^ | CT_DCE vs CT_Veh | 8 | Post Hoc Tests Multiple Comparisons Bonferroni | 0.0188 / * |
|  |  | CT_DCE vs CT_WT | 8 |  | 1 / N.S. |
|  |  | CT_Veh VS CT_WT | 8 |  | 0.0003 / * |
|  | NCAM1^32-35^ | CT_DCE vs CT_Veh | 8 | Post Hoc Tests Multiple Comparisons Bonferroni | 0.0257 / * |
|  |  | CT_DCE vs CT_WT | 8 |  | 0.7329 / N.S. |
|  |  | CT_Veh VS CT_WT | 8 |  | <0.0001 / * |
|  | PI4KA^36^ | CT_DCE vs CT_Veh | 8 | Post Hoc Tests Multiple Comparisons Bonferroni | 0.0035 / * |
|  |  | CT_DCE vs CT_WT | 8 |  | 1 / N.S. |
|  |  | CT_Veh VS CT_WT | 8 |  | 0.0263 / * |
|  | DCLK1^37^ | CT_DCE vs CT_Veh | 8 | Post Hoc Tests Multiple Comparisons Bonferroni | <0.0001 / * |
|  |  | CT_DCE vs CT_WT | 8 |  | 0.3225 / N.S. |
|  |  | CT_Veh VS CT_WT | 8 |  | <0.0001 / * |
|  | PURB | CT_DCE vs CT_Veh | 8 | Post Hoc Tests Multiple Comparisons Bonferroni | 0.0162 / * |
|  |  | CT_DCE vs CT_WT | 8 |  | 1 / N.S. |
|  |  | CT_Veh VS CT_WT | 8 |  | 0.0457 / * |

**Supplemental Table 3b**

| **Brain Region** | **Protein** | **Group Comparison** | **N number** | **Statistics** | **p value / Significance** |
| --- | --- | --- | --- | --- | --- |
| **Hippocampus** | NCAM1^31-35^ | Hippo_DCE vs Hippo_Veh | 8 | Post Hoc Tests Multiple Comparisons Bonferroni | 0.0041 / * |
|  |  | Hippo_DCE vs Hippo_WT | 8 |  | 1 / N.S. |
|  |  | Hippo_Veh VS Hippo_WT | 8 |  | 0.0383 / * |
|  | IGSF8^38^ | Hippo_DCE vs Hippo_Veh | 8 | Post Hoc Tests Multiple Comparisons Bonferroni | 0.0002 / * |
|  |  | Hippo_DCE vs Hippo_WT | 8 |  | 1 / N.S. |
|  |  | Hippo_Veh VS Hippo_WT | 8 |  | 0.0004 / * |
|  | KIF1A^39-42^ | Hippo_DCE vs Hippo_Veh | 8 | Post Hoc Tests Multiple Comparisons Bonferroni | <0.0001 / * |
|  |  | Hippo_DCE vs Hippo_WT | 8 |  | 0.0004 / * |
|  |  | Hippo_Veh VS Hippo_WT | 8 |  | <0.0001 / * |
|  | L1CAM^43^ | Hippo_DCE vs Hippo_Veh | 8 | Post Hoc Tests Multiple Comparisons Bonferroni | 0.0032 / * |
|  |  | Hippo_DCE vs Hippo_WT | 8 |  | 1 / N.S. |
|  |  | Hippo_Veh VS Hippo_WT | 8 |  | 0.0017 / * |

**Supplemental Table 3c**

| **Brain Region** | **Protein** | **Group Comparison** | **N number** | **Statistics** | **p value / Significance** |
| --- | --- | --- | --- | --- | --- |
| **Cerebellum** | LMNB1^44,45^ | CB_DCE vs CB_Veh | 8 | Post Hoc Tests Multiple Comparisons Bonferroni | <0.0001 / * |
|  |  | CB_DCE vs CB_WT | 8 |  | 1 / N.S. |
|  |  | CB_Veh VS CB_WT | 8 |  | <0.0001 / * |
|  | MYO5A | CB_DCE vs CB_Veh | 8 | Post Hoc Tests Multiple Comparisons Bonferroni | 0.0049 / * |
|  |  | CB_DCE vs CB_WT | 8 |  | 1 / N.S. |
|  |  | CB_Veh VS CB_WT | 8 |  | 0.4556 / N.S. |
|  | FABP7^46^ | CB_DCE vs CB_Veh | 8 | Post Hoc Tests Multiple Comparisons Bonferroni | <0.0001 / * |
|  |  | CB_DCE vs CB_WT | 8 |  | 1 / N.S. |
|  |  | CB_Veh VS CB_WT | 8 |  | <0.0001 / * |
|  | PURB | CB_DCE vs CB_Veh | 8 | Post Hoc Tests Multiple Comparisons Bonferroni | 0.0325 / * |
|  |  | CB_DCE vs CB_WT | 8 |  | 1 / N.S. |
|  |  | CB_Veh VS CB_WT | 8 |  | 0.0325 / * |
|  | MECP2^47-49^ | CB_DCE vs CB_Veh | 8 | Post Hoc Tests Multiple Comparisons Bonferroni | <0.0001 / * |
|  |  | CB_DCE vs CB_WT | 8 |  | 1 / N.S. |
|  |  | CB_Veh VS CB_WT | 8 |  | 0.0216 / * |
|  | ANK1 | CB_DCE vs CB_Veh | 8 | Post Hoc Tests Multiple Comparisons Bonferroni | <0.0001 / * |
|  |  | CB_DCE vs CB_WT | 8 |  | 0.846 / N.S. |
|  |  | CB_Veh VS CB_WT | 8 |  | 0.0002 / * |
|  | ANXA5^50^ | CB_DCE vs CB_Veh | 8 | Post Hoc Tests Multiple Comparisons Bonferroni | 0.0052 / * |
|  |  | CB_DCE vs CB_WT | 8 |  | 1 / N.S. |
|  |  | CB_Veh VS CB_WT | 8 |  | 0.0004 / * |
