## Supplemental Table 5 for "Dodecyl Creatine Ester Improves Cognitive Function and Identifies Drivers of Creatine Deficiency"

**Supplemental Table 5a: Proteins showing a significant correlation with cognitive performance in discrimination index test**

| **Proteins** | **Brain region** | **Correlation with cognitive performance (r^2^)** | **p value** |
| --- | --- | --- | --- |
| **KIF1A** | **Hippocampus** | **-0.442** | **0.015** |
| NCAM1 | Hippocampus | -0.361 | 0.042 |
| **LMNB1** | **Hippocampus** | **-0.634** | **<0.0001** |
| **L1CAM** | **Hippocampus** | **-0.367** | **0.039** |
| **FABP7** | **Hippocampus** | **0.548** | **0.003** |
| PLCB1 | Hippocampus | 0.460 | 0.012 |
| **Pi4KA** | **Hippocampus** | **-0.634** | **0.066** |

| **Proteins** | **Brain region** | **Correlation with cognitive performance (r^2^)** | **p value** |
| --- | --- | --- | --- |
| **KIF1A** | **Cortex** | **-0.506** | **0.006** |
| IGSF8 | Cortex | 0.411 | 0.023 |
| NCAM1 | Cortex | -0.567 | 0.002 |
| DCLK1 | Cortex | -0.431 | 0.018 |
| **L1CAM** | **Cortex** | **-0.376** | **0.035** |
| **FABP7** | **Cortex** | **-0.342** | **0.051** |
| Pi4KA | Cortex | -0.342 | 0.002 |
| ANXA5 | Cortex | 0.570 | 0.047 |

| **Proteins** | **Brain region** | **Correlation with cognitive performance (r^2^)** | **p value** |
| --- | --- | --- | --- |
| **KIF1A** | **Cerebellum** | **-0.466** | **0.011** |
| DCLK1 | Cerebellum | -0.362 | 0.041 |
| **L1CAM** | **Cerebellum** | **-0.433** | **0.017** |
| **FABP7** | **Cerebellum** | **-0.424** | **0.019** |
| PURB | Cerebellum | -0.274 | 0.038 |
| ANK1 | Cerebellum | 0.495 | 0.007 |

| **Proteins** | **Brain region** | **Correlation with cognitive performance (r^2^)** | **p value** |
| --- | --- | --- | --- |
| **KIF1A** | **Brain Stem** | **-0.366** | **0.040** |
| DCLK1 | Brain Stem | -0.584 | 0.001 |
| **L1CAM** | **Brain Stem** | **-0.464** | **0.001** |
| **FABP7** | **Brain Stem** | **-0.466** | **0.001** |
| ANK1 | Brain Stem | 0.400 | 0.026 |
| ANXA5 | Brain Stem | 0.480 | 0.009 |

**Supplemental Table 5b: Proteins showing a significant correlation with cognitive performance in Y-maze test**

| **Proteins** | **Brain region** | **Correlation with cognitive performance (r^2^)** | **p value** |
| --- | --- | --- | --- |
| **KIF1A** | **Hippocampus** | **-0.795** | **<0.0001** |
| NCAM1 | Hippocampus | -0.375 | 0.036 |
| **L1CAM** | **Hippocampus** | **-0.386** | **0.050** |
| **IGSF8** | **Hippocampus** | **-0.344** | **0.003** |
| MYO5 | Hippocampus | -0.450 | 0.014 |
| ANXA5 | Hippocampus | 0.523 | 0.004 |

| **Proteins** | **Brain region** | **Correlation with cognitive performance (r^2^)** | **p value** |
| --- | --- | --- | --- |
| **KIF1A** | **Cortex** | **-0.608** | **0.001** |
| **IGSF8** | **Cortex** | **0.439** | **0.016** |
| **NCAM1** | **Cortex** | **-0.494** | **0.007** |
| DCLK1 | Cortex | -0.566 | 0.002 |
| **L1CAM** | **Cortex** | **-0.580** | **0.001** |
| **FABP7** | **Cortex** | **-0.411** | **0.023** |
| PLCB1 | Cortex | 0.469 | 0.010 |
| PURB | Cortex | -0.363 | 0.041 |

| **Proteins** | **Brain region** | **Correlation with cognitive performance (r^2^)** | **p value** |
| --- | --- | --- | --- |
| **KIF1A** | **Cerebellum** | **-0.774** | **<0.0001** |
| **IGSF8** | **Cerebellum** | **0.374** | **0.035** |
| **L1CAM** | **Cerebellum** | **-0.431** | **0.018** |
| **FABP7** | **Cerebellum** | **-0.395** | **0.028** |
| LMNB1 | Cerebellum | -0.381 | 0.033 |
| ANXA5 | Cerebellum | 0.569 | 0.002 |

| **Proteins** | **Brain region** | **Correlation with cognitive performance (r^2^)** | **p value** |
| --- | --- | --- | --- |
| **FABP7** | **Brain Stem** | **-0.345** | **0.049** |
| LMNB1 | Brain Stem | -0.429 | 0.018 |
